# A Comparative Assessment of Visceral Leishmaniasis Burden in Two Eco-epidemiologically Different Countries, India and Sudan

**DOI:** 10.1101/592220

**Authors:** Kamal Barley, Anuj Mubayi, Muntaser Safan, Carlos Castillo-Chavez

## Abstract

The two hyper–endemic regions for Visceral Leishmaniasis (VL) in the world are located in India and Sudan. These two countries account for more than half of the world’s VL burden. The regional risk factors associated with VL vary drastically per region. A mathematical model of VL transmission dynamics is introduced and parametrized to quantify risk of VL infection in India and Sudan via a careful analysis of VL prevalence level and the control reproductive number, 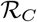, a metric often used to characterize the degree of endemicity. Parameters, associated with VL-epidemiology for India and Sudan, are estimated using data from health departmental reports, clinical trials, field studies, and surveys in order to assess potential differences between the hyper–endemic regions of India and Sudan. The estimated value of reproduction number for India is found to be 60% higher than that of Sudan (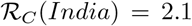 and 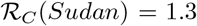). It is observed that the 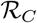 is most sensitive to the average biting rate and vector-human transmission rates irrespective of regional differences. The treatment rate is found to be the most sensitive parameter to VL prevalence in humans for both India and Sudan. Although the unexplained higher incidence of VL in India needs to be carefully monitored during long-term empirical follow-up, the risk factors associated with vectors are identified as more critical to dynamics of VL than factors related to humans through this modeling study.

**Author Summary:** The Visceral Leishmaniasis (VL) is a neglected tropical disease, primarily endemic in five countries, with India and Sudan having the highest burden. The risk factors associated with VL are either unknown in some regions or vary drastically among empirical studies. In this study, we collect VL-related data from multiple sources for the two different countries, India and Sudan, and use techniques from mathematical modeling to understand factors that may be critical in the spread and control of VL. The results suggest that the risk factors associated with disease progression are important in explaining high VL prevalence in both the countries. However, the likelihood of disease outbreak in India is much higher than that in Sudan and the probability of transmission between human and sandfly populations vary significantly between the two. The results have implications towards VL elimination and may require a review of current control priorities.

## 1 Introduction

### Leishmaniasis Globally

Leishmaniasis is a family of infectious diseases caused by an intracellular protozoan parasite of the genus *Leishmania* [80]. A diverse and complex pathogen, Leishmania can be transmitted to humans through the bite of one of at least 20 different species of female sand flies of the subfamily Phlebotomus [17,42]. Individuals living with Leishmaniasis may exhibit one of the four clinical syndromes; cutaneous, mucocutaneous, diffuse cutaneous, and visceral Leishmaniasis. [17, 77]. Visceral Leishmaniasis (VL, also known as Kala-Azar (KA) in Hindi) is considered the most severe form of the disease because death is inevitable if untreated. In fact, there are significant distinctions that have been observed even in the dynamics of VL from one region to another. VL is most often caused by species of the *Leishmania donovani* complex with *Leishmania donovani sensu sticto* circulating in the Indian subcontinent, *Leishamania donovani sensu lato* in East Africa, *Leishmania infantum* primarily found around the Mediterranean, in the Middle East, and rest of the Africa and Asia and *Leishmania chagasi* in the Americas [70]. There are marked differences between parasite species infections, for example, in terms of epidemiology, clinical features and responses to treatment.

### Epidemiology of VL

*Leishmania donovani* (*L. donovani*) infects VL in most affected regions, with each year there is an estimated 500,000 new cases and approximately 50,000 recorded deaths worldwide [19]. Researchers estimate that roughly 12 million people are infected with *Leishmania* parasites, at a given time, among the 350 million individuals at risk [4,61]. However, these statistics might be changing with recent WHO’s efforts in eliminating VL from some parts of the world. VL is endemic in at least 88 tropical and subtropical countries around the world with more than 90% of new cases generated in Bangladesh, Brazil, India, Nepal, and Sudan [32, 74]. In 2010, the state of Bihar in India reported an average of 270,000 new cases per year with an incidence rate of 21 cases per 1000 [42]. Twenty one districts out of Bihar’s 38 districts are most affected from VL. The most recent (2014) report estimates that there are between 200,000 and 400,000 annual cases of VL in the five most affected countries, with India supporting between 146,700 to 282,000 cases per year and Sudan between 15,700 and 30,300 cases per year [9]. In Sudan, VL is endemic in southern, central, and eastern parts of the country, with most cases being reported from state of Gedaref (near the Ethiopian border) [49]. VL primarily affects low socio-economic and marginalized communities [57]. Geographic hot spots for infection are characterized by factors that include the average length of the sand flies life cycle, the abundance of parasite reservoirs, and human behaviors to infection [32, 74]. In this study, we aim to identify factors associated with VL burden in the two most affected countries in the world, India and Sudan.

### Risk Factors of VL in India and Sudan

In India, the sand fly species *Phlebotomus Argentipes* is primarily responsible for transmitting the *L. donovani* parasite [67]. In Indian state of Bihar, annual patterns of VL incidence are assumed to be driven by ecological and social factors including distinct seasonality in sand fly population, lack of health care resources, extreme poverty, frequent flooding resulting in food shortages, and malnutrition [6, 57]. In Sudan, *Phlebotomus Orientalis* is the dominant sandfly vector associated with anthroponotic *L. donovani* transmission. [28, 31, 38, 39, 72, 86]. Typically, *P. Orientalis* is considered a forest species and its abundance is frequently associated with the presence of the savanna woodland tree species Acacia Seyal and Balanites aegyptiaca and deeply cracked vertisols (black cotton soil) [28, 29]. Primary risk factors for VL infection in Sudan include genetic factors (e.g., some indigenous individuals may be more susceptible [6]), age, ethnicity, the consequences of poverty, movements of people facing civil war, and political instability which is accompanied by labour migrations for economic security reasons [6, 15, 68].

### Interventions in India and Sudan

In Bihar, where 90% of India’s VL cases occur, aggressive attempts at improving vector control programs via the distribution of insecticide-treated bed nets and insecticide spraying are being carried out [6]. India’s Kala-azar Elimination Programs (KAEP) aims at reducing VL morbidity are tied into government-funded VL diagnosis and drug treatment programs. Pentavalent antimonial drugs, wherever it is effective, purchased by the public sector are barely sufficient to cover half of the infected patients [5, 57]. Limited drug availability and drug resistance are growing problems in East Africa, particularly in Sudan, where antimonials are still the primary method of VL medical treatment. The poor must travel long distances to gain access to drugs and, consequently, the effectiveness of intervention policies are limited. Infected Sudanese often must wait extended periods of time before receiving minimal medical care [62].

### VL Mathematical Modeling Studies

In 1988, Dye and Wolpert introduced what it appears to be the first Anthroponotic VL deterministic model for capturing the temporal dynamics of this disease. Their model was used to explain the observed VL inter-epidemic periods between 1875 and 1950 in Assam, India. Following this work, Dye, C. (1992, 1996) assessed the impact of control measures on VL patterns in endemic areas using appropriately modified models [23–25]. These studies concluded that dramatic upswing in VL cases in the past may be attributed to “intrinsic” factors related primarily to disease epidemiology in humans and vectors and not to “extrinsic” processes. Mathematical models are typically developed to capture VL transmission dynamics in a population. Using such model, a threshold quantity for infection, the reproductive number 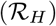, is often computed for understanding the dynamics of the disease. 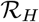 is used to measure the disease’s ability to colonize a naive population or to identify the degree of endemicity in the presence of intervention. In general, model analysis suggests disease persistence when 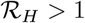 and eventual disease extinction when 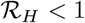 [84].

### Focus of this Study

In the hyper–endemic regions of India and Sudan, it is believed that 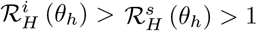, where 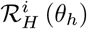, the control reproductive number, for India is greater than 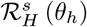 of Sudan at their respective rate of treatment, *θ_h_*. This modeling study focuses exclusively on the transmission dynamics and control of VL in India and Sudan while proving or refuting the belief on differences in their estimated reproduction numbers. Since VL is hyper–endemic in these regions for long time, a model with established treatment regimes is considered the “status quo” an important assumption, since untreated individuals die relatively quickly. Parameters (transmission rates, death rates, etc.) are estimated using novel simple methods where current treatment (*θ_h_*) is always present. Consequently, the process of invasion (ability of VL to invade a population) is addressed under current treatment policies. Therefore, the reproductive number includes treatment rate, *θ_h_*, as part of the initial set-up where detailed infection data is absent (technically it cannot be called the “basic” reproductive number). A comparative study of the VL situation in India and Sudan is carried out via a model derived metrics parameterized using estimates derived from published clinical trials data and published national reports. Uncertainty and sensitivity analyses are then carried out to identify key risk factors and use them to evaluate the effectiveness of intervention programs in the two “worst-affected” VL-regions of the world. In summary, the goals of this study are: (i) identify relevant data from field/clinical studies needed to estimate model parameters of a dynamic model, (ii) develop procedures to estimate distributions of quantities for which data are unavailable, (iii) evaluate risk associated with VL in India and Sudan, and (iv) compare and contrast the risk factors between India and Sudan. The details of the analysis are depicted in the Figure 1.

**Figure 1.**
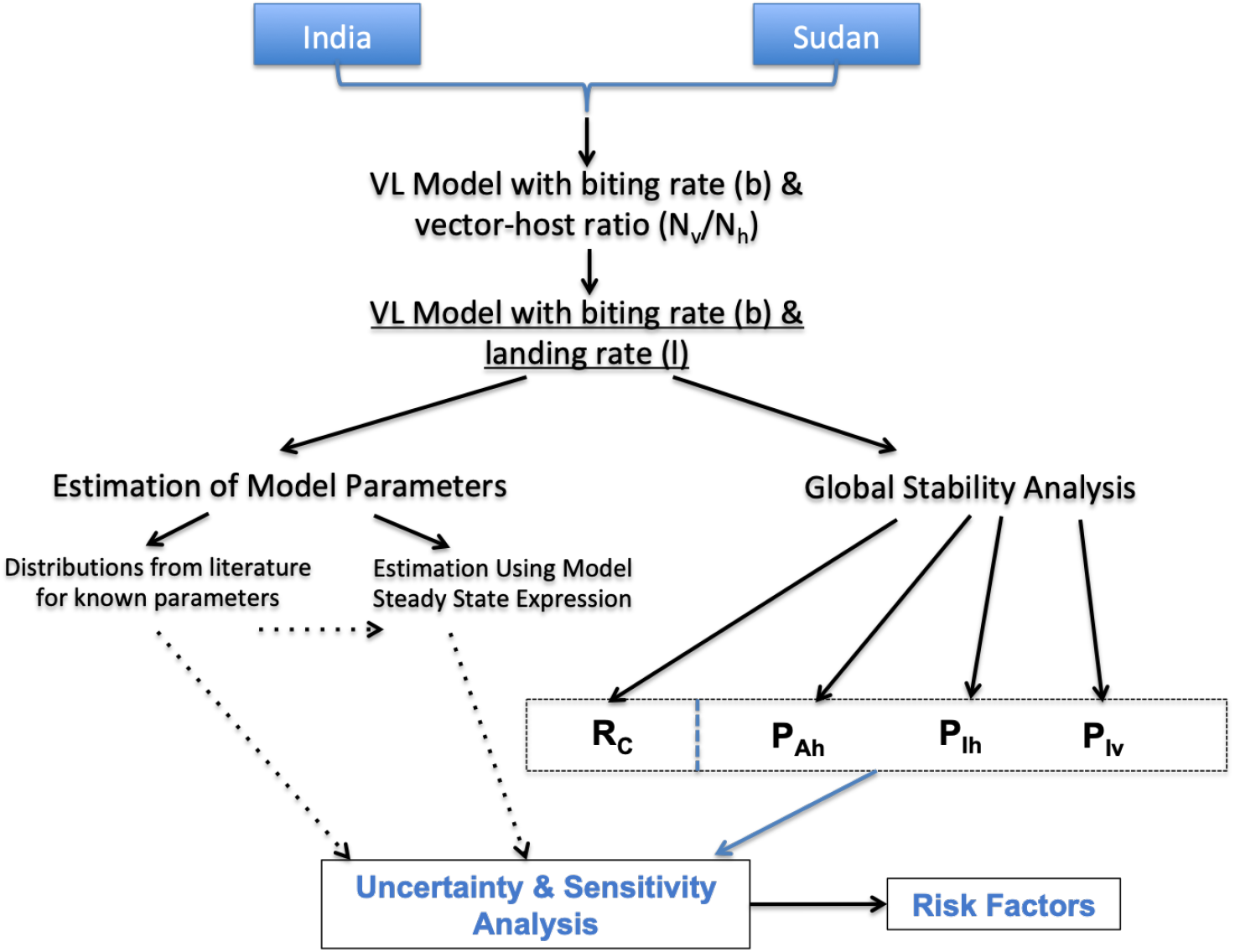
Flow chart representing steps in the analysis.

## 2 Methods

### 2.1 Model formulation and assumptions

The *Leishmania donovani* transmission cycle is anthroponotic and takes place from human to human via the bite of an infective female phlebotomine Sandfly. A mathematical model of the transmission dynamics of VL infection is used here where the interacting host (*N_h_*(*t*)) and vector (*N_v_*(*t*)) populations are assumed to mix homogeneously. The flow chart representing disease progression and transmission is shown in Figure 2. The human population is subdivided into susceptible individuals (*S_h_*(*t*)), asymptomatic individuals (*A_h_*(*t*)), infectious individuals with clinical VL infection (*I_h_*(*t*)), individuals under treatment (*T_h_*(*t*)), and recovered-immune to reinfection individuals (*R_h_*(*t*)); *N_h_* ≡ *S_h_* + *A_h_* + *I_h_* + *T_h_* + *R_h_*. The sandfly population is assumed to be divided into susceptible (*S_v_*(*t*)) and infectious (*I_v_*(*t*)) vectors with *N_v_* ≡ *S_v_* + *I_v_*.

**Figure 2.**
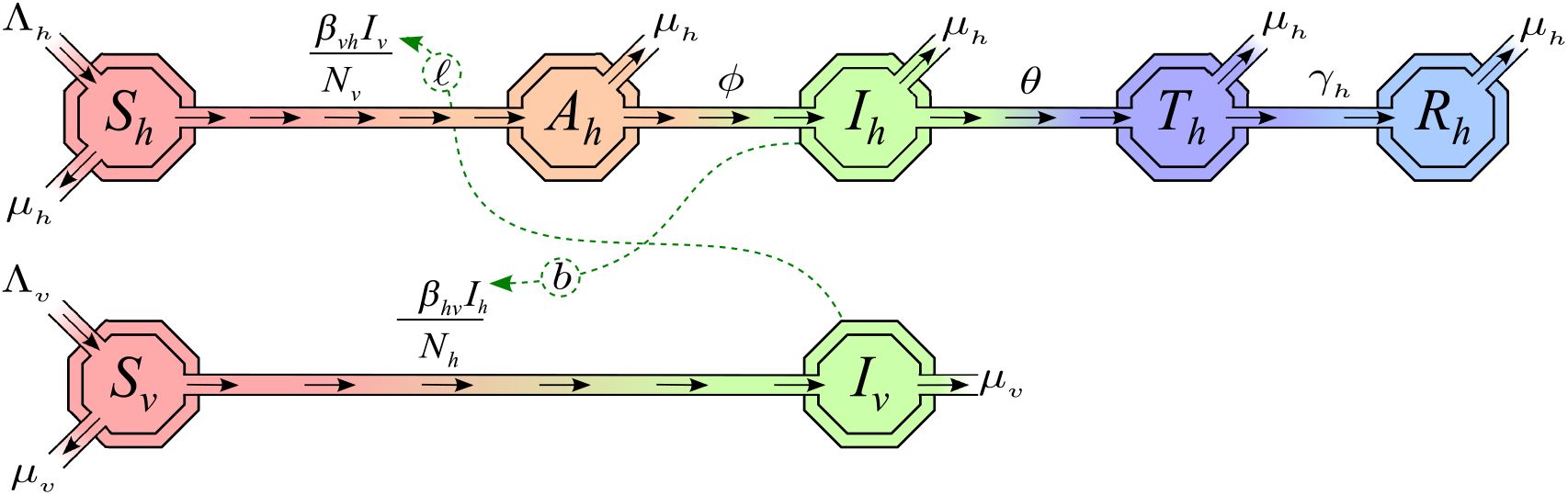
A schematic representation of the mathematical modeling framework consisting of interacting human (*N_h_*) and Sandfly (*N_v_*) populations. Arrows represent transition between different infection stages in the two populations.

The model system is given as:

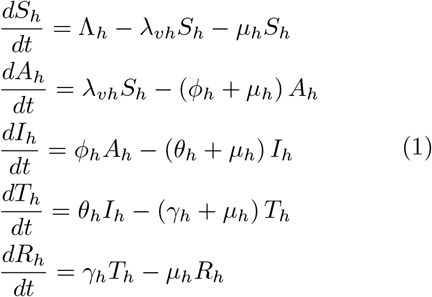

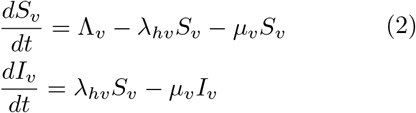

**Table 1.**
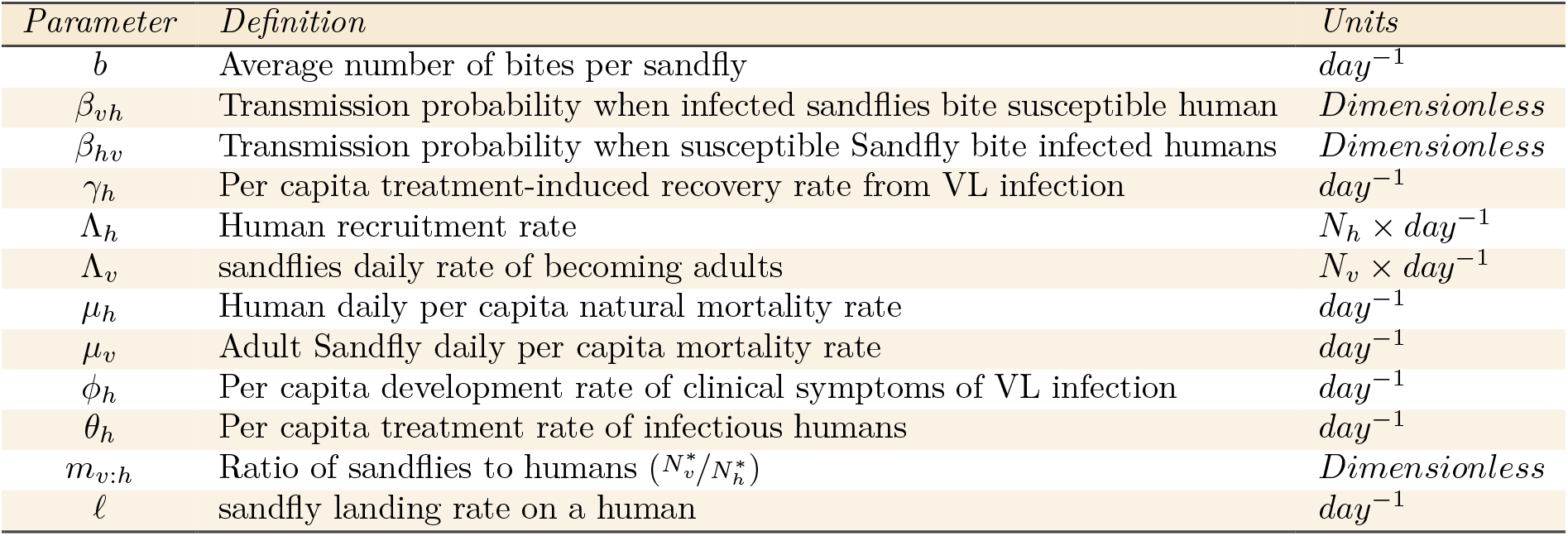
The parameters for the VL model and their dimensions

Disease-induced mortality is not included because, due to institutionalized treatment, deaths from VL are negligible. For simplicity, the human population is assumed to be constant. Λ_*h*_ denotes the recruitment rate into the susceptible population, and *μ_h_* denotes the per-capita death rate. Because *N_h_* approaches 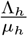 when *t* approaches ∞, we assume, without loss of generality, that 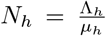 [16]. A susceptible individual acquires the *L. Donovani* parasite following an effective contact with an infectious sandfly. The rate *λ_vh_*, the force of infection on humans, is given by

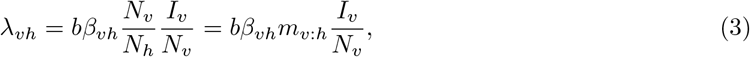

where the right-hand expression (Equation 3) is given by the product of the per-vector daily biting rate of sandflies (*b*), the VL infection transmission probability, given a bite from an infected sandfly to human (*β_vh_*), the average number of sandflies per humans *m_v:h_*, and the proportion of infectious sandflies in the vector population (I_v_/N_v_). It is assumed that all newly VL-infected humans go through an asymptomatic (symptomless) stage (*A_h_*). After an asymptomatic period of several months, humans develop clinical symptoms at the per capita rate *ϕ_h_*, moving to the infectious class *I_h_*. During the infectious period, humans will seek VL treatment at the per capita rate *θ_h_*, proper treatment leads to recovery at the per capita rate *γ_h_* (recovered individuals gain lifelong immunity). Newly emerging adult female sandflies are recruited into the susceptible population at rate Λ_*v*_ and die at the per-capita rate *μ_v_*. The sandfly population is assumed constant. A susceptible sandfly is infected following an effective contact with infectious humans at the per capita rate *λ_hv_* (force of infection on sandflies). The rate *λ_hv_* is given by

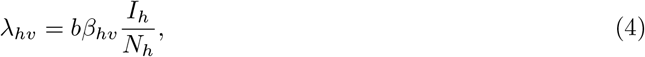

where the right-hand side is the product of: the per vector daily biting rate (*b*); the probability that susceptible sandflies acquire the *Leishmania* parasite while feeding on a VL-infected individuals (*β_hv_*); the proportion of VL infectious humans in the human population (I_h_/N_h_,). It is also assumed that the *Leishmania* parasite has no impact on an infected sandfly’s lifespan; the sandflies’ natural mortality percapita rate is the same for infected and uninfected, namely, *μ_v_*. See Appendix B for complete model derivation.

### 2.2 Biting rates

Interactions between vector biting behavior and uneven pathogen transmission potential between hosts may lead to difficulty in controlling infection. How vector species respond to availability of hosts is highly variable and has fostered considerable interest among vector borne disease modelers for decades. The proportion of blood-meals taken by vectors from the host species of interest is generally assumed to increase directly with increasing human availability and changing levels of vector density. Hence, vector biting can play a significant role in the transmission process [88]. The biting rate of sandflies is typically a function of ambient air temperatures, humidity, wind speed, vector density and local habitat. There remains a couple of challenges in effectively using biting rates, namely, which is a proper functional response to capture biting rates in the model and how to measure it precisely from the field data.

To effectively use models to make reasonable definitions, models must be carefully parameterized and validated with epidemiological and entomological data. On the other hand, researchers have modeled biting rate in different ways but realistically the biting rate may vary according to the abundance of hosts and to vector preference [87]. In this study, we suggest alternative forms of transmission terms as well as use distinct data sets to estimate parameters of the two different terms (vector-to-host and host-to-vector terms).

### 2.3 Incidence as a Function of the Landing Rates

This section first provides a careful derivation of incidence rates expression as a function of landing and biting rates and then use landing rate data to estimate the transmission probabilities from sandflies to humans (*β_vh_*) and humans to sandflies (*β_hv_*).

The human incidence rate (Equation (3)) is a function of the average rate of interactions between vectors and humans, which in turn is directly proportional to the proportion of infectious sandflies 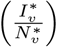. Let *b* denote the average number of bites per sandfly per unit time and *ρ* the average number of bites received per human per unit time. *Assuming that all sandfly bites are to humans only, we must have that the total number of bites made by all sandflies per unit of time* 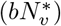 *equals the total number of bites received by all human hosts per unit of time* 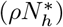. Thus, we have that

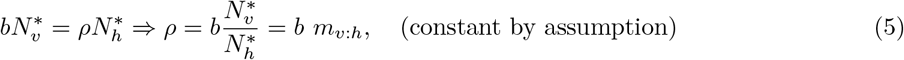

The assumption that *ρ* is constant is customary in the literature although there are some studies where the host vector ratio is assumed not constant over time [85]. *We further assume that the average number of bites received by a human per unit time is proportional to the number of sandflies landing on an individual per unit time, that is*,

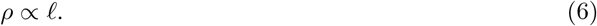

Hence, the total number of effective landings on all humans from all sandflies per unit time is

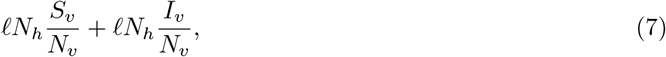

where the first (second) term of (7) accounts for the total number of effective landings on all humans from all susceptible (infected) sandflies per unit time. That is, the total effective landing/feeding of vectors on humans is a function of the total vector population, which includes both susceptible and infected vectors. In epidemiology, of importance are only the two cases when landing occurs from a susceptible sandfly on an infected human and from an infected sandfly on a susceptible human, as they are the cases where landing results in transmission of VL from humans to sandflies and vice versa. In other words, *ℓN_h_* is the total number of effective landings per unit time, while 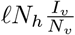 is the proportion of bites that result in infecting new hosts. Therefore, 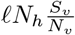 is the proportion of bites that get “wasted” since they cannot generate infections.

If *β_vh_* is the per-person transmission efficiency (that is, probability that infection is successfully transmitted from vector to human given an infected bite), then the rate at which VL is transmitted to humans is

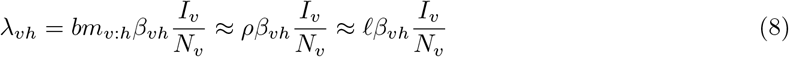

using Equations (5) and (6).

Similarly, we can derive the infection rate in the vector population generated by infected humans. If 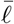 accounts for the average number of times a sandfly lands on humans per unit time, then the total number of effective landings by all sandflies on all humans is

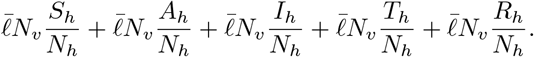

It should be noted that the total 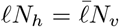and that, while accounting for new incidences in sandflies, we are interested in landings occurring from susceptible sandflies on infected humans only. Hence, the term

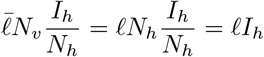

is the one that plays a role in accounting for new sandflies incidences, while the remaining terms aren’t. If we let *β_hv_* be the per-person transmission efficiency from human to vector (i.e., transmission probability per bite on infectious humans that leads to infection in a susceptible sandfly), then the total number of sandflies who acquire infection while effectively landing on infected humans per unit time is

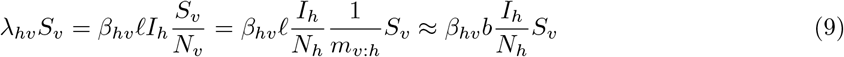

using Equations (5) and (6).

## 3 Analysis

In this section, we derive from the model an expression for the average number of secondary infections generated by an infected individual (referred here as the control reproduction number), as well as expressions for the prevalence of different types of the populations. We also discuss the procedures used for estimating model parameters.

### 3.1 Stability Analysis

The analysis of Model (1)–(2) shows that it has two equilibriums, namely, the Disease Free Equilibrium (DFE) and Endemic Equilibrium (EE). The existence and stability of the equilibriums depends on the threshold ratio, the reproduction number, first introduced by Sir Ronald A. Ross in his 1911 seminal work on malaria [41] and it provides a measure of the risk posed by an invading disease in a population without any intervention for the disease. The control reproduction number, 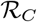, is a similar ratio defined as the number of secondary infections caused by a single infective introduced in a primarily susceptible population (i.e., *N* ≈ *S*_0_) but in the presence of interventions [12, 20, 53, 84].

Using our model, we compute 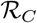 using the next generation operator approach [12, 13, 84], a process that requires the computation of the matrix of new infection terms, **F**, and the matrix of transition between compartments, **V**. The 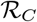 is the spectral radius of the next generation matrix, *ρ* (**FV**^−1^) (see section B.1 for derivation), in the presence of treatment program (where the treatment rate is *θ_h_*) and is given by

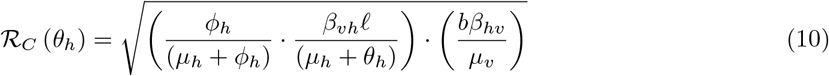

where *ℓ* is the landing rate on a human, *b* is the biting rate per sandfly, and *β_vh_* the number of infections in humans generated by one infected vector. The expression 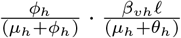 is the average number of new cases vectors generated by one infected human and 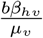 represent average number of new cases in humans produced by one infected vector. Hence, 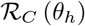 is given by the geometric mean of two sub “reproduction” numbers

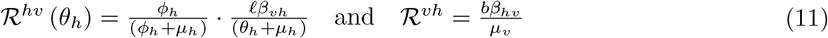

where 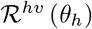 is interpreted as the number of secondary infections caused in humans through a bite of a single typical infectious sand fly into an entirely susceptible host population in the presence of treatment program while 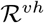 denotes the number of secondary infections in female sandflies caused by one newly introduced infected human.

*Remark* 3.1. The DFE of Model (1–2) always exist and is globally asymptotically stable (LAS) if 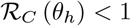 and unstable if 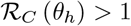. (see Appendix B.4 for proof)

*Remark* 3.2. The EE exists and is globally stable only when 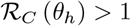 (see Section B.4 for proof).

### 3.2 Robustness Analysis

VL has received comparatively much less attention by researchers and policy makers as compared to many other tropical diseases and hence, is classified as one of the neglected diseases by WHO. There are limited number of studies that collect data to study VL patterns and even fewer studies that use such data in a dynamical model for evaluating control programs. In this research, we carry out a thorough literature review to identify what data is available and what is missing that may be needed to understand comprehensively VL dynamics for two most affected countries in the world.

Since 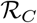 plays a key role in the transmission dynamics of VL and parameters are often not precisely measured in India and Sudan, studying parameter sensitivity of the model outputs including on 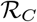 becomes important if we wish to identify the pressure points of the system. Uncertainty (UA) and sensitivity (SA) analyses are used here to assess the robustness in the model results as a function of uncertainty in the estimated model parameters from available data. The analyses rely on the Latin Hypercube Sampling (LHS) and require the computation of the Partial Rank Correlation Coefficient (PRCC), a sensitivity index with respect to each of the model parameters [11, 54, 55]. The LHS scheme includes the generation of a stratified random sampling that ensures a systematic optimal exploration of the feasible parameter space. In the sampling, an input parameter *X* with a pre-defined probability distribution function (PDF) is divided into *N* equiprobable subintervals. From each subintervals, a value is sampled. The *N* values for this parameter are randomly paired with the the corresponding *N* values of other parameters generated in the same way. The PRCCs is used to measure the degree of linear association between a model output and a parameter from a set of parameters, after influence of linearity from all other parameters of the set had been eliminated [54]. The calculated PRCCs and corresponding p-values value are used to rank sensitivity of the parameters to the output variable. The PRCC value of each imput parameter is considered statistically significant, with *p*-value< 0.05, if PRCC > |0.3|.

Multiple data sources and reports were considered to obtain point estimates for each of the model parameters for which precise value was not obtained. Using the point estimates, a theoretical distribution is fitted to available data for corresponding parameter and random samples were generated via distribution. We assess the impact of variation in model parameters on estimates, as well as the level of influence, of each, on estimates of 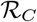 and the country specific prevalence. We develop approaches using our dynamic model to estimate country specific (India and Sudan) parameters for which data was unavailable and performed parametric uncertainty and sensitivity analysis on model based metrics that defines risk based on four different definitions: (i) 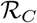, (ii) prevalence of asymptomatic humans, (iii) prevalence of symptomatic humans, or (iv) prevalence of infectious vectors.

## 4 Results

### 4.1 Parameter Estimation

Model parameter estimates, and their ranges, and distributions were obtained for India and Sudan using prevalence data, published literature, and methodology developed in sections below [22,27,51,77,86]. In the case of the species of *Phlebotomus sandflies*, most of the parameter estimates were taken from data collected via field studies in parasitology and ecology literature [22, 27, 51]. We provide details of all parameter estimates in Section C of Appendix and a summarize them in Table 2. We estimated parameters, for which precise data could not be obtained for India and Sudan, via our two developed approaches. In the next Section 4.1.2 we give a detailed discussion and procedure for estimating transmission probabilities of the model for both countries.

**Table 2.**
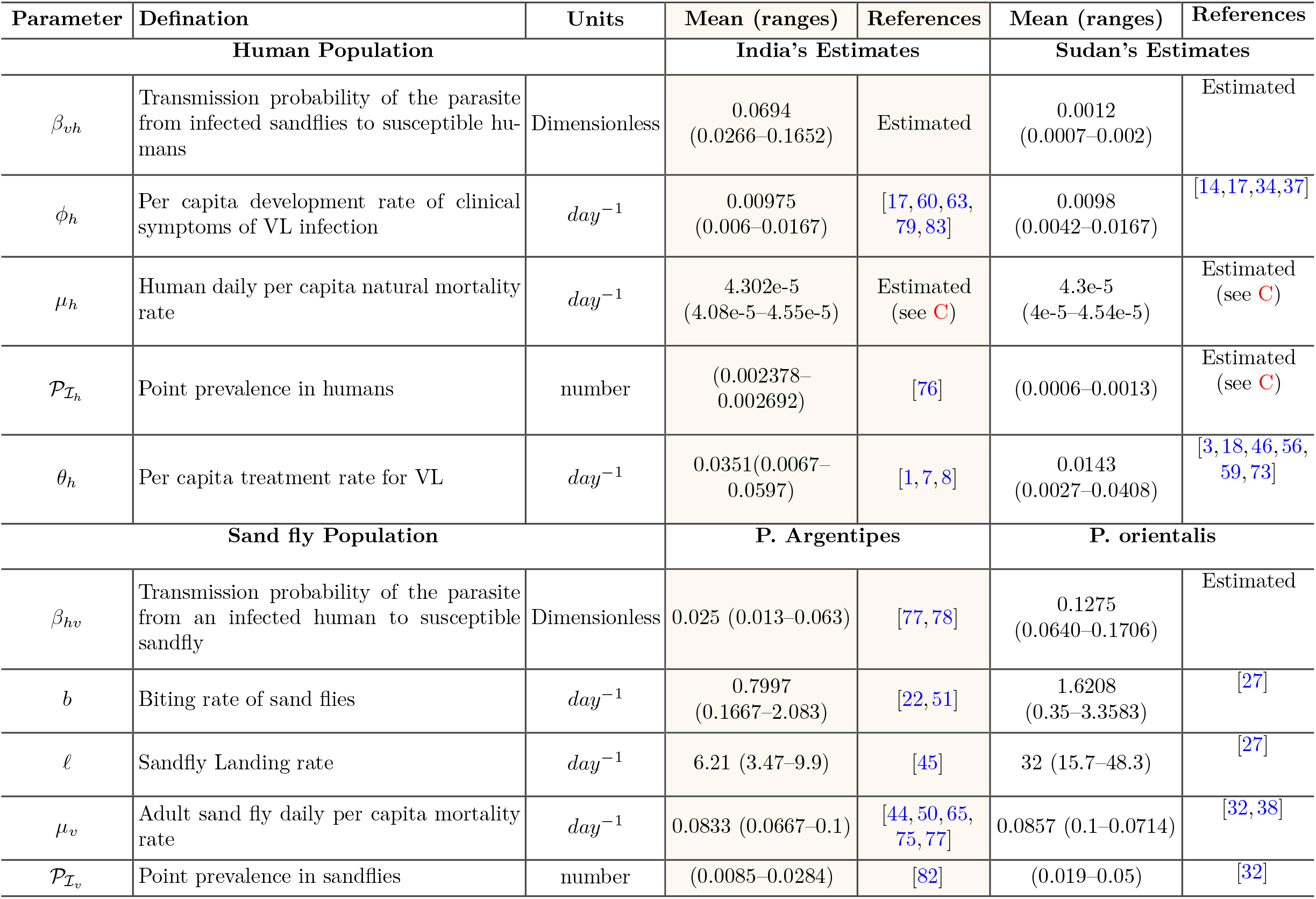
Model parameter estimates related to human and vector (Phlebotomus sand fly species) populations in India and Sudan

#### 4.1.1 Landing rate

The nocturnal activities of various sandfly species start at around 6:00 pm – 9:00 pm, peaks between the hours of 11:00 pm – 1:00 am and ends between the hours of 3:00 pm and 6:00 am. A rapid rise to a maximum pick and then a sharp decline observed in data from various field studies suggest the probability distribution for biting and landing rate would best be fitted with a triangular distribution. However, most data represented more closely to landing rates and hence, in this section we show the fits of landing rate distribution. From the fitted data for each respective countries, the shape parameters (min, max, and mode) for the triangular distribution for landing rate was estimated from the sandflies trap data (see Figures 3a and 3b for *P. Argentines* and Figures 4a and 4b for *P. Orientalis*).

**Figure 3.**
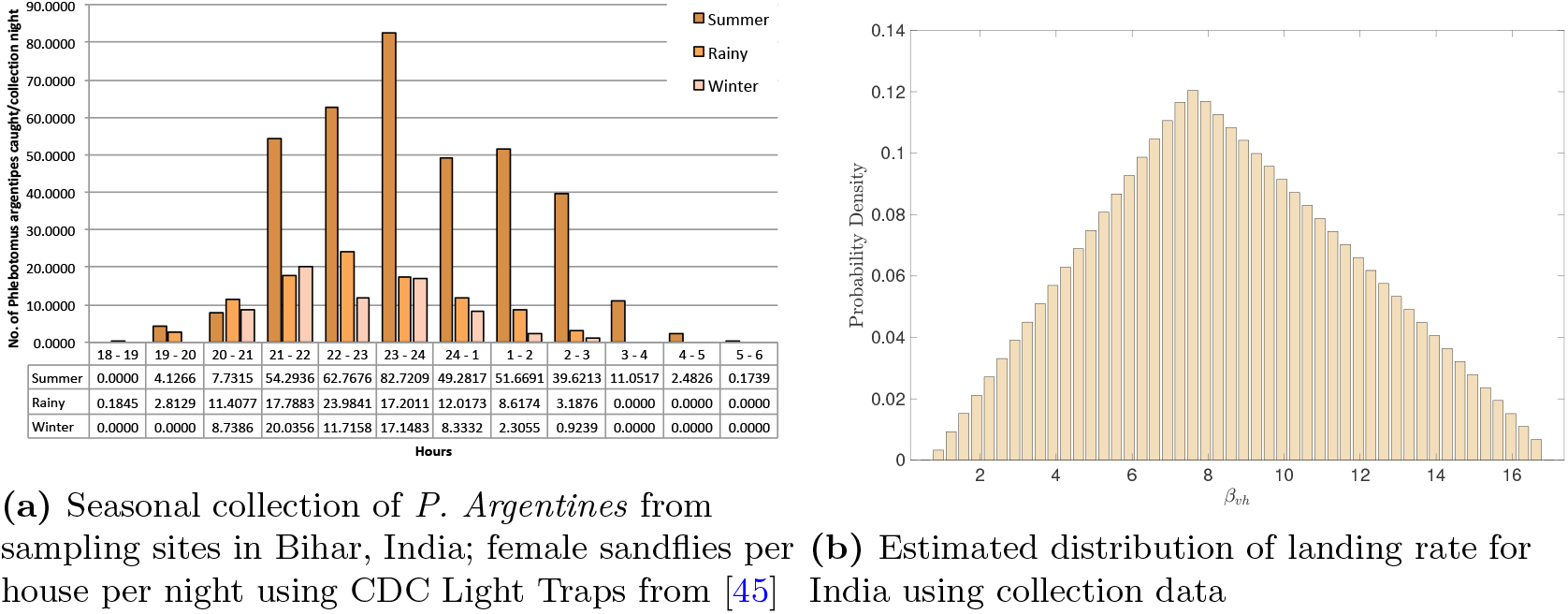
Collected data of *P. Argentines* was first averaged over seasons and then fitted to the triangular distribution to estimate parameters of the distribution representing landing rate. The mean and 95% Confidence Interval for landing rate distribution are given in Table 2.

**Figure 4.**
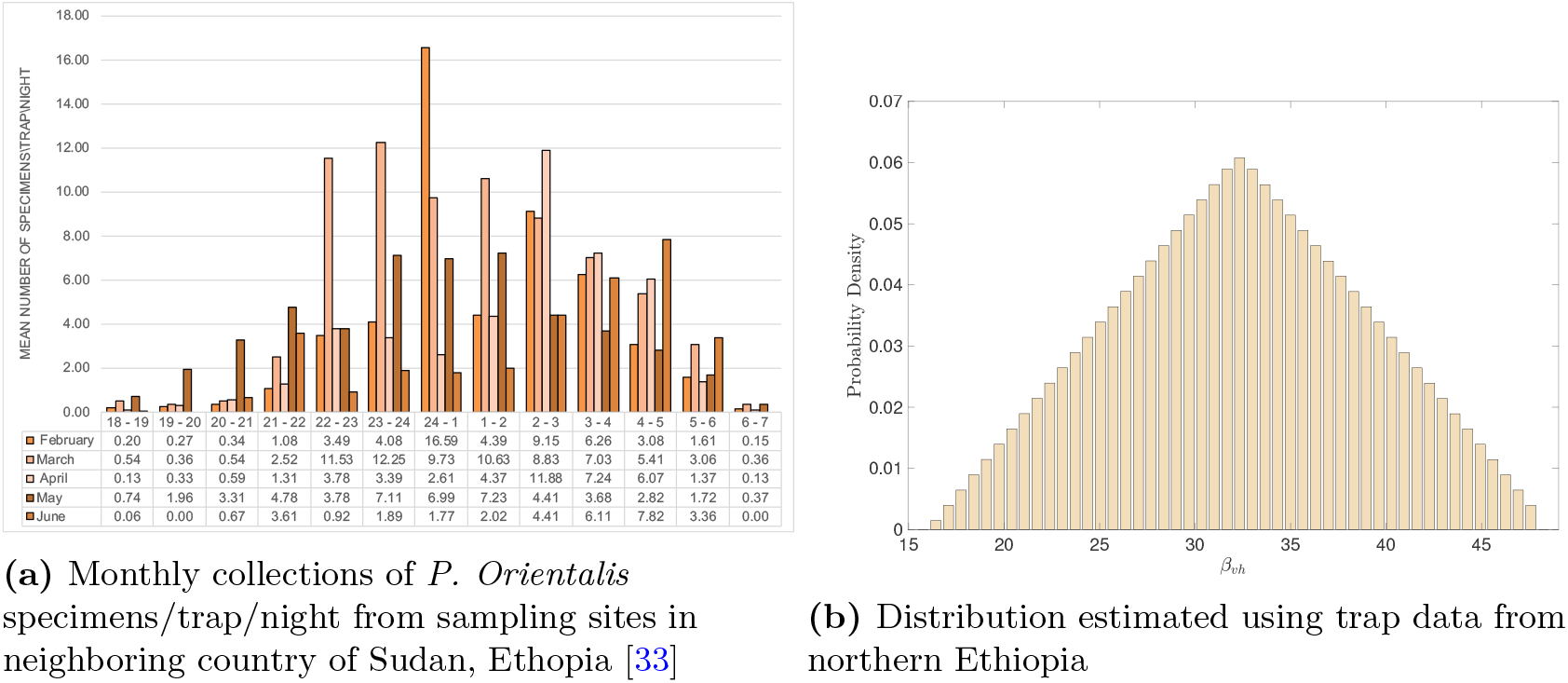
Collected data of *P. Orientalis* was first averaged over months and then fitted to the triangular distribution to estimate parameters of the distribution representing landing rate. The mean and 95% Confidence Interval for landing rate distribution are given in Table 2.

#### 4.1.2 Approaches for Estimating the Transmission Probabilities

Lack of active surveillance, effective case identification and case management results in under-reporting of cases and uncertainties in epidemiological parameter. A survey of the literature on mathematical studies on VL dynamics revealed that estimates obtained for the transmission probabilities for VL are often based on corresponding estimates for malaria, dengue and other well-studied vector-borne diseases. Consequently, borrowing of parameter estimates from other established vector-borne models can contribute epistemic uncertainties in the epidemic threshold and underestimate or overestimate model predictions. To understand the impact of these uncertainties on parameter estimates, we used ranges for parameters for which we can obtain data with relatively high certainty and mathematical methods to estimate the parameters representing transmission probabilities required in our model. Two novel approaches, that uses endemic prevalence from the model, were designed to estimate the transmission probabilities as described in the next two sub-sections. Note, the unique endemic equilibrium of the model is stable (as shown in the Section 3) and is given by

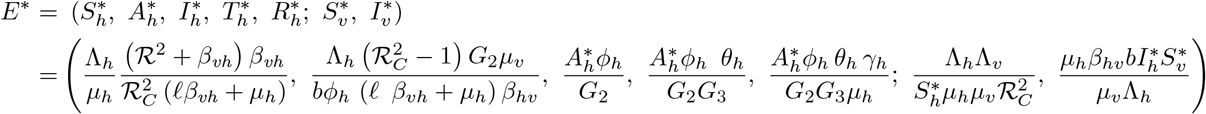

where *G*_1_ = *ϕ_h_* + *μ_h_, G*_2_ = *θ_h_* + *μ_h_*, and *G*_3_ = *γ_h_* + *μ_h_*. The explicit expressions of the infected components of the endemic equilibrium are

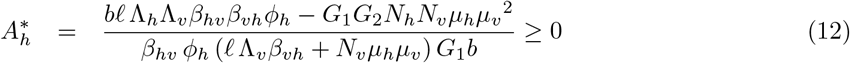

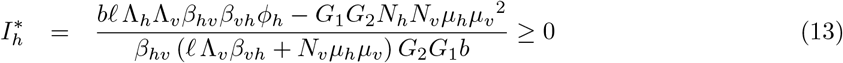

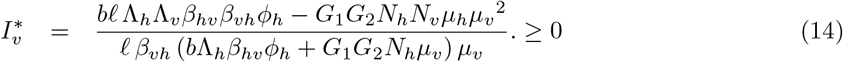

Since VL is endemic in both India and Sudan, we use these expression to obtain prevalences and thereby use them to estimate transmission probabilities (i.e. *β_vh_, β_vh_*). We assume Λ_*h*_ = *μ_h_N_h_* and Λ_*v*_ = *μ_v_N_v_* and hence, the host and vector populations becomes constant. The prevalences in humans and sandflies populations are given by

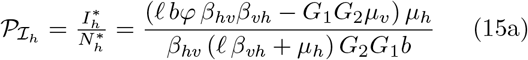

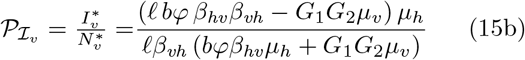

##### Approach 1

Fixing all model parameters for which data was available and assuming that humans and vectors prevalences are known, we obtain simultaneous equations in *β_hv_* and *β_vh_* using Equations (15a) and (15b). Solving the simultaneous equations, we get

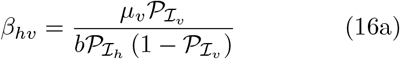

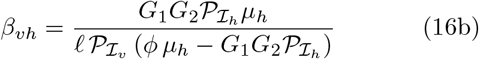

The equations (16a) and (16b) along with the estimates of other model parameters and known sample host and vector prevalences are used to obtain estimates of the transmission probabilities. The estimated distributions using the this approach are given in Figure 5.

**Figure 5.**
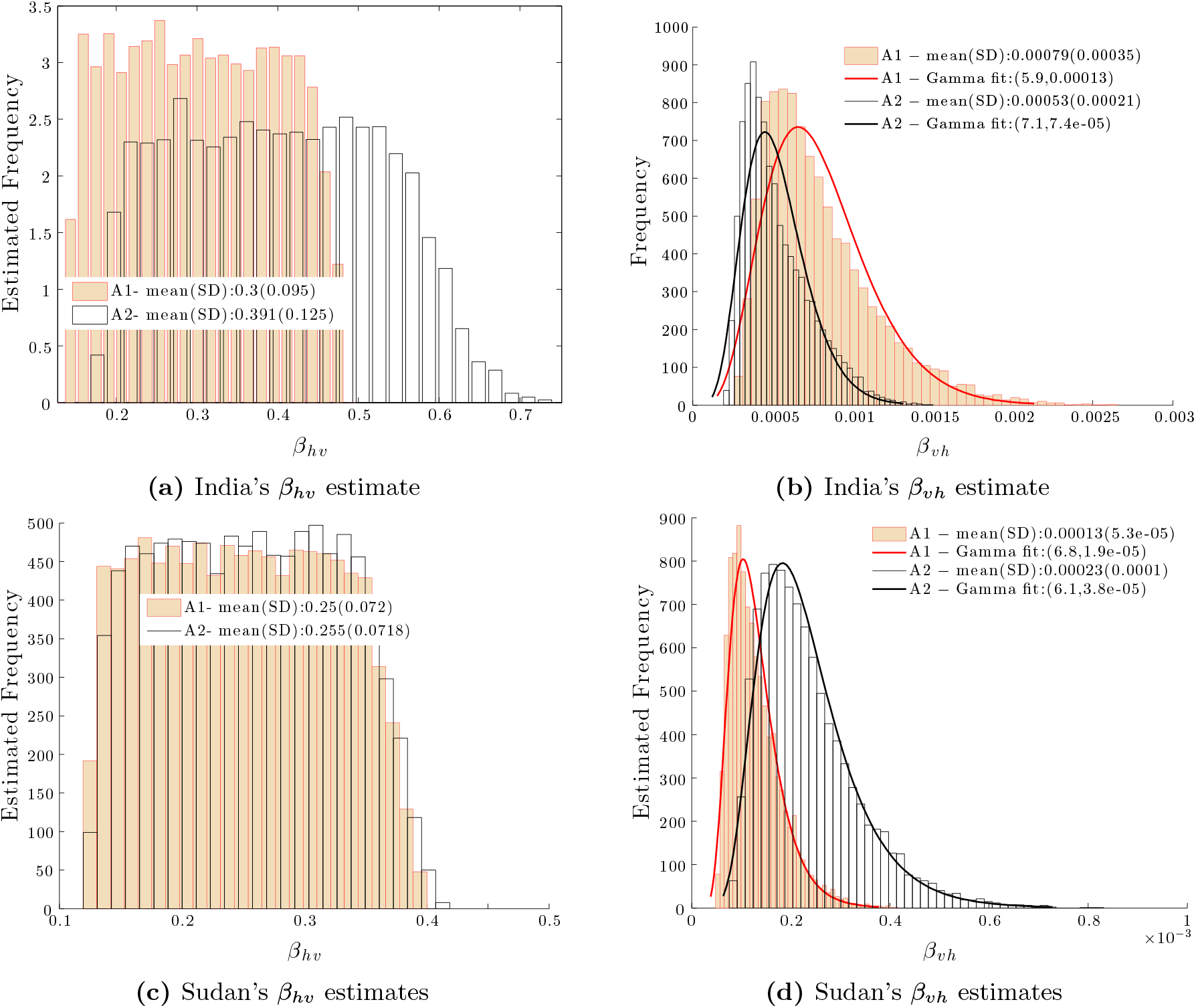
Estimated distribution of *β_vh_* and *β_hv_* for India (a–b) and Sudan (c–d), respectively. A1 (A2) represents the distribution obtained using Approach 1 (Approach 2). A visual comparison of the fitted gamma distribution together with the model obtained estimated transmission probabilities, *β_vh_*.

##### Approach 2

In this approach, we rely on estimates of 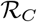 from modeling studies in literature to estimate the transmission probabilities for the two countries. Using Equation (10), the expressions (15a) and (15b) for the prevalences can be rewritten in terms of 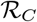 as follows:

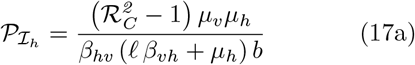

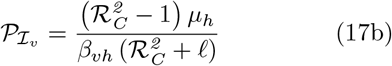

Isolating *β_vh_* and *β_hv_* from (17a) and (17b) we obtain,

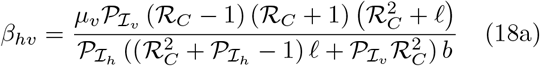

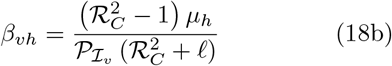

The estimated distributions using the this approach are given in Figure 5.

### 4.2 Parameter uncertainty and sensitivity analyses

Parameter uncertainty and sensitivity analyses are performed on two different quantities: the reproduction number (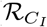 for India and 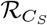 for Sudan) and the Prevalence of the infected populations (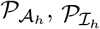, and 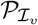). These analyses are used to assess which of the eight input parameters (*b, ℓ, β_hv_, β_vh_, μ_h_, μ_v_, ϕ_h_*, and *θ_h_*) are most significant to estimating disease patterns.

#### 4.2.1 On 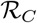

Uncertainty and sensitivity analyses on the control reproduction number 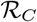 (the outcome variable) assess critical parameters to disease dynamics. We fit a parametric probability density function (PDF) for each of the eight parameters to respective available data. In the case of the parameters *b, μ_v_, m_h:v_, β_vh_, β_hv_, μ_h_*, and *μ_v_* a uniform distribution was generated since the minimum and maximum point estimates were only found in the literature. The parameters, *ϕ_h_*, and *θ_h_* were assigned a gamma distribution as estimated in previous study based on inverse problem approach (Mubayi et al. [57]). For each of the eight parameters with assigned probability distributions, sample sizes of 10, 000 values were randomly generated over ten independent realizations. Using LHS technique, in each of the realizations we paired randomly the first *N* samples of the first column (samples of first parameter) with *N* samples from the second column (samples of second parameter). After all eight parameters were paired without replacement; an LHS matrix was generated with rows and columns corresponding to parameters and entries of the LHS samples. Each row of parameters in the LHS matrix were considered to be random inputs variables for generating one value of 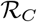 using Equation (10). Thus, *N* × *p* LHS matrix (where *p* represents number of parameters on which 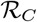 depends) results in *N* samples for 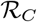 in each realization.

After 10 realizations, the mean (*μ*) of point estimates of 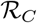, standard error (*σ*) of 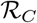, and the probabilities that 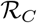 estimates fall below and above the threshold value one for India and Sudan were collected (Table ??). The mean estimated value for 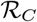 for India was found to be approximately 2.11, which greater than the corresponding mean estimated value of 1.30 for Sudan. The fact that India has the highest estimated incidence in the world (146,700 to 282,800 per year) roughly twice of that in Sudan having the highest in Africa (15,700 to 30,300 per year) [5,6], is not enough unless we are able to re-scale them appropriately, to make any conclusions on that largest differences in 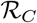-values. These estimates of 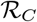 confirm the current VL status in India and Sudan. The difference in the magnitude of 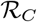 may be attributable to the fact that India carries a much greater burden of all new VL cases (almost more than 50%) worldwide.

Statistical analysis on the differences in the means of 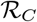 for India and Sudan was carried out using a t-test with *H*_0_: *M_I_* = *M_S_* against *H*_1_: *M_I_* ≠ *M_S_* where the mean of 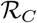 for India was denoted as *M_I_* and for Sudan *M_S_*. The analysis suggested rejection of null hypothesis (Table 4), that is, the obtained point estimates of 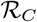 between India and Sudan are statistically different. Now that we have concluded that 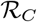 for India is indeed significantly higher than that the one for Sudan using model generated 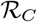, re-scaled by population size, we proceed to determine what are the parameters that if modified generates the larger change in 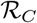.

**Table 3.** Summary of estimates of the transmission probabilities, *β_hv_* and *β_vh_*, using the two approaches with mean and ranges for other parameters (Table 2) for India and Sudan were fixed.

|  | India |  |  | Sudan |  |  |
| --- | --- | --- | --- | --- | --- | --- |
| Parameters | Min | Mean | Max | Min | Mean | Max |
| **Fixed** |  |  |  |  |  |  |
| $b$ | - | 2.08 | - | - | 1.6208 | - |
| $\mu_h$ | - | 4.54e-5 | - | - | 4.3e-5 | - |
| $\mu_v$ | - | 0.0833 | - | - | 0.0857 | - |
| $\phi_h$ | - | 0.00975 | - | - | 0.0098 | - |
| $\theta_h$ | - | 0.0083 | - | - | 0.0143 | - |
| **Varied** |  |  |  |  |  |  |
| $\mathcal{P}_h$ | 0.0024 | - | 0.0027 | 0.0013 | - | 0.0015 |
| $\mathcal{P}_v$ | 0.0054 | - | 0.0157 | 0.054 | - | 0.037 |
| $\ell$ | 8.68 | 12.15 | 17 | 15.7 | 32 | 48.3 |
| $\beta_{hv}$ | 0.025 | 0.012 | 0.038 | 0.0032 | 0.0167 | 0.041 |
| $\mathcal{R}_C$ | 1.3 | 2.0 | 2.1 | 1.1 | 1.3 | 1.5 |
| **Estimates using Approach 1** |  |  |  |  |  |  |
| $\beta_{vh}$ | 0.21 | 0.44 | 0.7 | 0.24 | 0.53 | 0.95 |
| $\beta_{hv}$ | 0.00015 | 0.00035 | 0.00091 | 0.00013 | 0.00042 | 0.0012 |
| **Estimates using Approach 2** |  |  |  |  |  |  |
| $\beta_{vh}$ | 0.19 | 0.4 | 0.64 | 0.2 | 0.41 | 0.68 |
| $\beta_{hv}$ | 4.9e-05 | 0.00013 | 0.0004 | 3.6e-05 | 0.00011 | 0.00037 |

**Table 4.** Mean *R_C_* estimates and results of statistical test for testing differences of *R_C_* between India and Sudan

| Country | $R_C$ estimated values | |
| --- | --- | --- |
| India | $M_I$: Mean(SD) | 2.11 (1.6) |
| Sudan | $M_S$: Mean(SD) | 1.31 (0.79) |
| | 95% CI of $\vert M_I - M_S\vert $ | (0.77, 0.84) |
| | T-test | $H_0 : \vert M_I - M_S\vert = 0$ |
| | | $P - value < 0.05$ |
|  | Test statistic | tstat(Value of the test statistic): 146.0191 |
|  |  | Degrees of freedom of the test : 19998 |

The PRCCs were calculated for each country in order to quantify sensitivity of model parameters on the 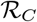 estimates. We observe the sign and the magnitude of the PRCC values for each parameter above the line *y* = ±0.3 for each respective country.

#### 4.2.2 On endemic prevalences (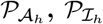, and 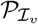)

Parameter uncertainty and sensitivity analyses are also performed on the Prevalence of the infected populations (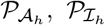, and 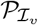). These analysis are used to assess which of the same eight input parameters (*b, ℓ, β_hv_, β_vh_, μ_h_, μ_v_, ϕ_h_*, and *θ_h_*) are most significant to estimating endemic prevalences.

As described in the Section 4.2.1 on *R_C_*, the similar sensitivity and uncertainty analysis procedure was carried out on the endemic prevalences for both the countries. However, higher number of samples (50, 000) for each parameter were obtained. The first 10, 000 sample-sets (out of the 50, 000) that resulted in 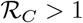 (condition for existence of the endemic prevalence) were eventually used in the analysis. This is because that endemic equilibrium only exists and stable when 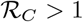.

### 4.3 Assessment for India

#### 4.3.1 Uncertainty and Sensitivity Analysis on 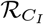

The estimated distribution of 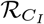 from uncertainty analysis, is shown in Figure 7a. The mean estimate of 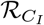 for India is found to be 2.05 with a standard deviation of 1.09. The sensitivity analysis of 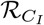 provides the ranking of parameters based on their influence on 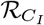 (Figure 7e). In decreasing order of influence, the parameter ranking was *θ_h_*, being the most sensitive parameter, followed by *b, ℓ, β_vh_, β_hv_*, and the least sensitive parameters are *ϕ_h_* followed by *μ_v_*.

**Figure 6.**
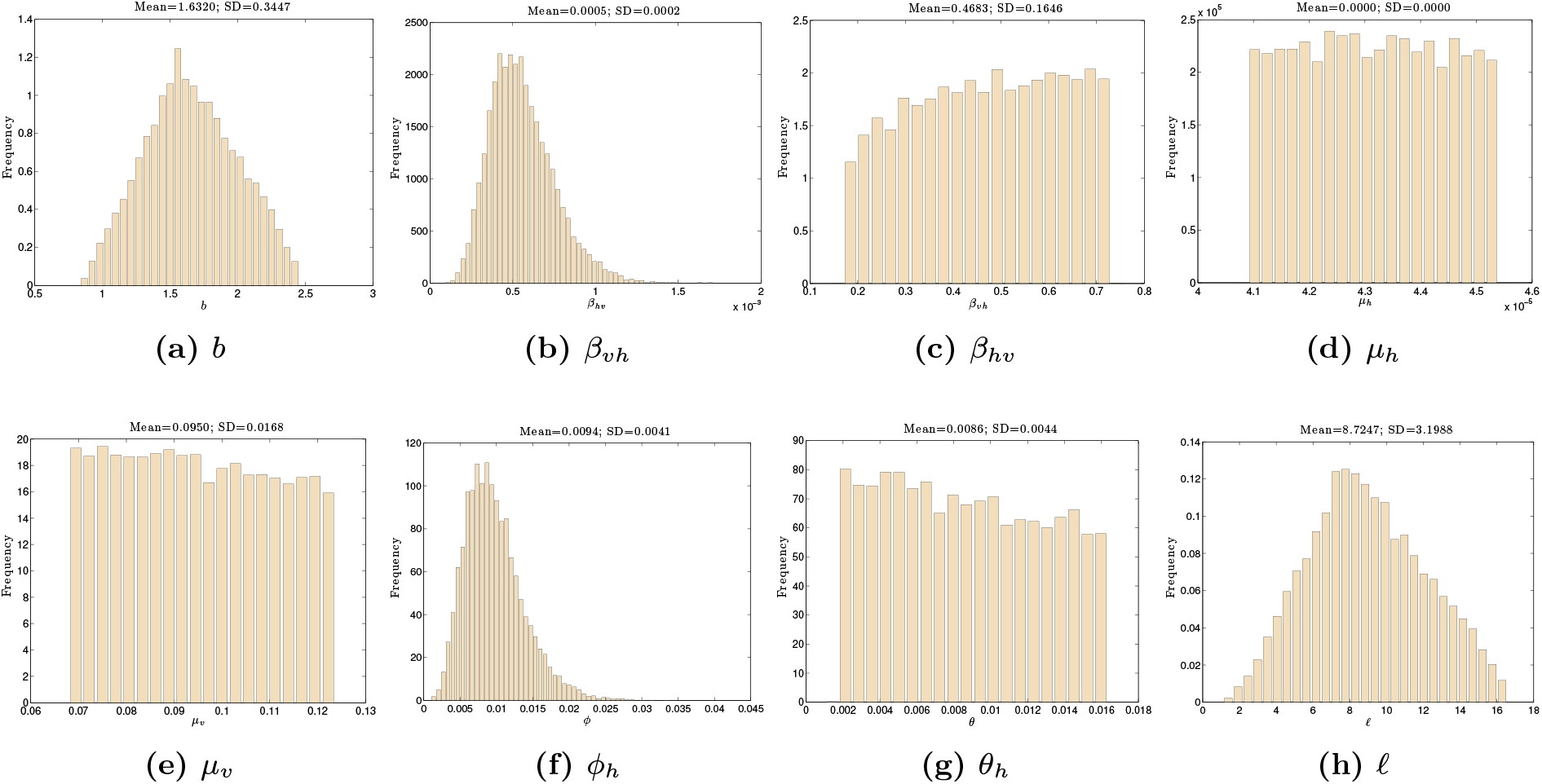
Parameter distributions conditional on 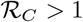 for India obtained from uncertainty analysis of the prevalence

**Figure 7.**
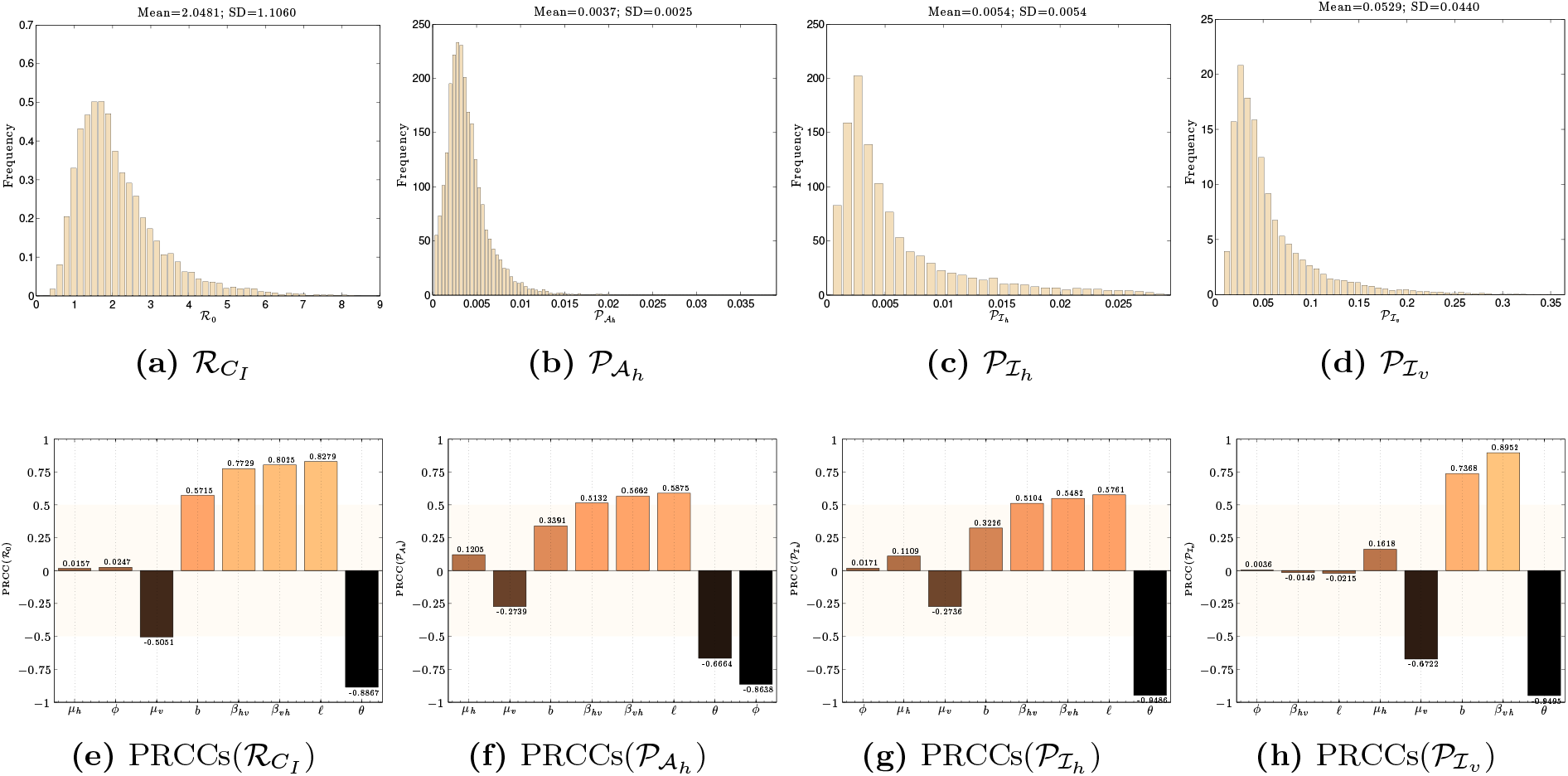
Results for India: Uncertainty in the Reproduction Number (Subfigure 7a) and the Prevalence (Subfigures 7b –7d) of Asymtomatics, Infectious Humans and Infectious Sandflies, respectively. Tornado plot showing partial rank correlation coefficients (PRCCs) of the Reproduction number (7e) and the Prevalence (7f –7h) in Asymtomatics, Infectious Humans and Infectious Sandflies, respectively.

#### 4.3.2 Uncertainty and Sensitivity Analysis on the three Endemic Prevalences

The estimated distributions of prevalence (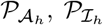, and 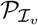) are shown in Figure 7b–7d. The mean estimate of 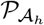 was found to be 0.0045 with a standard deviation of 0.0019. The parameter *ϕ_h_* was found to be the most influential parameter on the prevalence of asymptomatic, 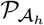. The remaining parameters in descending order of magnitude of PRCC were, *θ_h_, ℓ, β_vh_*, and *β_hv_*, with *μ_v_* and *μ_h_* being least sensitive parameters to 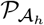. The sensitivity analysis performed on 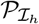 reveal that the treatment rate, *θ_h_* is the most influential parameter for changing disease prevalence. The mean estimates of vector prevalence were found to be 0.0526 with a with a standard deviation of 0.0432. From our sensitivity analysis of 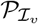 we observe in Table 7 and Figure 7h, that there are four most influential parameters. These parameters in decreasing order of ranks, are *θ_h_, β_vh_, b* and *μ_v_*.

**Table 5.** Comparing and contrasting risk factors that are critical to VL dynamics between India and Sudan.

| Similarity & Differences |  |
| --- | --- |
| Critical risk factors for the VL dynamics in both countries are: (i) Sandfly biting rates and (ii) Transmission rates between vector and human host | Treatment rate is critical for controlling outbreaks in India but less important in case of controlling outbreaks in Sudan. |
| *Infection transmission related (mean estimates for India are higher than that of Sudan)* |  |
| **India** [mean (std)]<br>$\beta_{vh} = 0.45$ (0.17)<br>$\beta_{hv} = 0.0005$ (0.0002)<br>$\mathcal{R}_C = 2.1$ (1.1)<br>$P(\mathcal{R}_C > 1) = 0.73$ | **Sudan** [mean (std)]<br>$\beta_{vh} = 0.27$ (0.09)<br>$\beta_{hv} = 0.0002$ (0.0001)<br>$\mathcal{R}_C = 1.3$ (0.6)<br>$P(\mathcal{R}_C > 1) = 0.58$ |
| *Sandfly ecology related (mean estimates for Sudan are higher than that of India)* |  |
| **India** [P. Argentipes]<br>$b = 1.6$ per day (Avg. biting rate)<br>$\ell = 8.3$ per day (Avg. landing rate)<br>$1/\mu_v = 10.0$ days (Avg. adult life span) | **Sudan** [P. Orientalis]<br>$b = 1.8$ per day<br>$\ell = 32.0$ per day<br>$1/\mu_v = 11.1$ days |

**Table 6.** Estimated parametric distribution of the model parameters for India and Sudan. The notations are → Triangular: 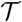(*min, mode, max*); Gamma: 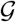(*shape, scale*); Uniform: 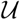(*min,max*).

| Parameter | India | Sudan |
| --- | --- | --- |
| $b$ | $\mathcal{T}(0.8, 1.6, 2.5)$ | $\mathcal{T}(0.35, 1.8, 3.4)$ |
| $\ell$ | $\mathcal{T}(0.55, 8.3, 17)$ | $\mathcal{T}(16, 32, 48)$ |
| $\phi_h$ | $\mathcal{G}(5.5470, 0.0021)$ | $\mathcal{G}(5.2727, 0.0018)$ |
| $\beta_{vh}$ | $\mathcal{G}(7, 7.5e-05)$ | $\mathcal{G}(6.3, 3.7e-05)$ |
| $\beta_{hv}$ | $\mathcal{U}(0.16, 0.73)$ | $\mathcal{U}(0.12, 0.42)$ |
| $\theta_h$ | $\mathcal{U}(0.0014, 0.0167)$ | $\mathcal{U}(0.0082, 0.0329)$ |
| $\mu_h$ | $\mathcal{U}(4.1e-5, 4.5e-5)$ | $\mathcal{U}(4e-5, 4.5e-5)$ |
| $\mu_v$ | $\mathcal{U}(0.0667, 0.1250)$ | $\mathcal{U}(0.071, 0.1)$ |

**Table 7.** Shows the PRCCs by rank of importance for the input parameters of the output value 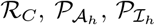, and 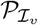 for India. (*) denotes PRCCs that are non-significant.

| Output | $\mathcal{R}_{C_I}$ | | $\mathcal{P}_{A_h}$ | | $\mathcal{P}_{I_h}$ | | $\mathcal{P}_{I_v}$ | |
| --- | --- | --- | --- | --- | --- | --- | --- | --- |
| Rank | Parameter | PRCC | Parameter | PRCC | Parameter | PRCC | Parameter | PRCC |
| 1 | $\theta_h$ | -0.89 | $\phi_h$ | -0.86 | $\theta_h$ | -0.95 | $\theta_h$ | -0.95 |
| 2 | $\ell$ | 0.83 | $\theta_h$ | -0.67 | $\ell$ | 0.58 | $\beta_{vh}$ | 0.9 |
| 3 | $\beta_{vh}$ | 0.8 | $\ell$ | 0.59 | $\beta_{vh}$ | 0.55 | $b$ | 0.74 |
| 4 | $\beta_{hv}$ | 0.77 | $\beta_{vh}$ | 0.57 | $\beta_{hv}$ | 0.51 | $\mu_v$ | -0.67 |
| 5 | $b$ | 0.57 | $\beta_{hv}$ | 0.51 | $b$ | 0.32 | $\mu_h$ | 0.16* |
| 6 | $\mu_v$ | -0.51 | $b$ | 0.34 | $\mu_v$ | -0.27* | $\ell$ | -0.021* |
| 7 | $\phi_h$ | 0.025* | $\mu_v$ | -0.27* | $\mu_h$ | 0.11* | $\beta_{hv}$ | -0.015* |
| 8 | $\mu_h$ | 0.016* | $\mu_h$ | 0.12* | $\phi_h$ | 0.017* | $\phi_h$ | 0.0036* |

### 4.4 Assessment for Sudan

#### 4.4.1 Uncertainty and Sensitivity Analysis on 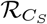

The result of uncertainty analysis on 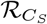 is shown in Figure 9a, where the mean estimate of 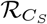 is 1.43, and the standard deviation is 0.6. From the Table 8 and Figure 9a we observe *β_hv_, b, θ_h_, β_vh_, ℓ*, and *μ_v_* are most sensitive (in order of ranking) to 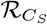. The first negatively correlated parameter was *θ_h_* which indicated that treatment is effective for controlling infection, followed by *μ_v_* which may relate to the impact of vector related control programs. The top two most positive parameters (i.e., positive PRCC) were *β_hv_* and *b*, which indicates that sandflies parameters may play a significant role in the estimation of 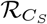.

**Figure 8.**
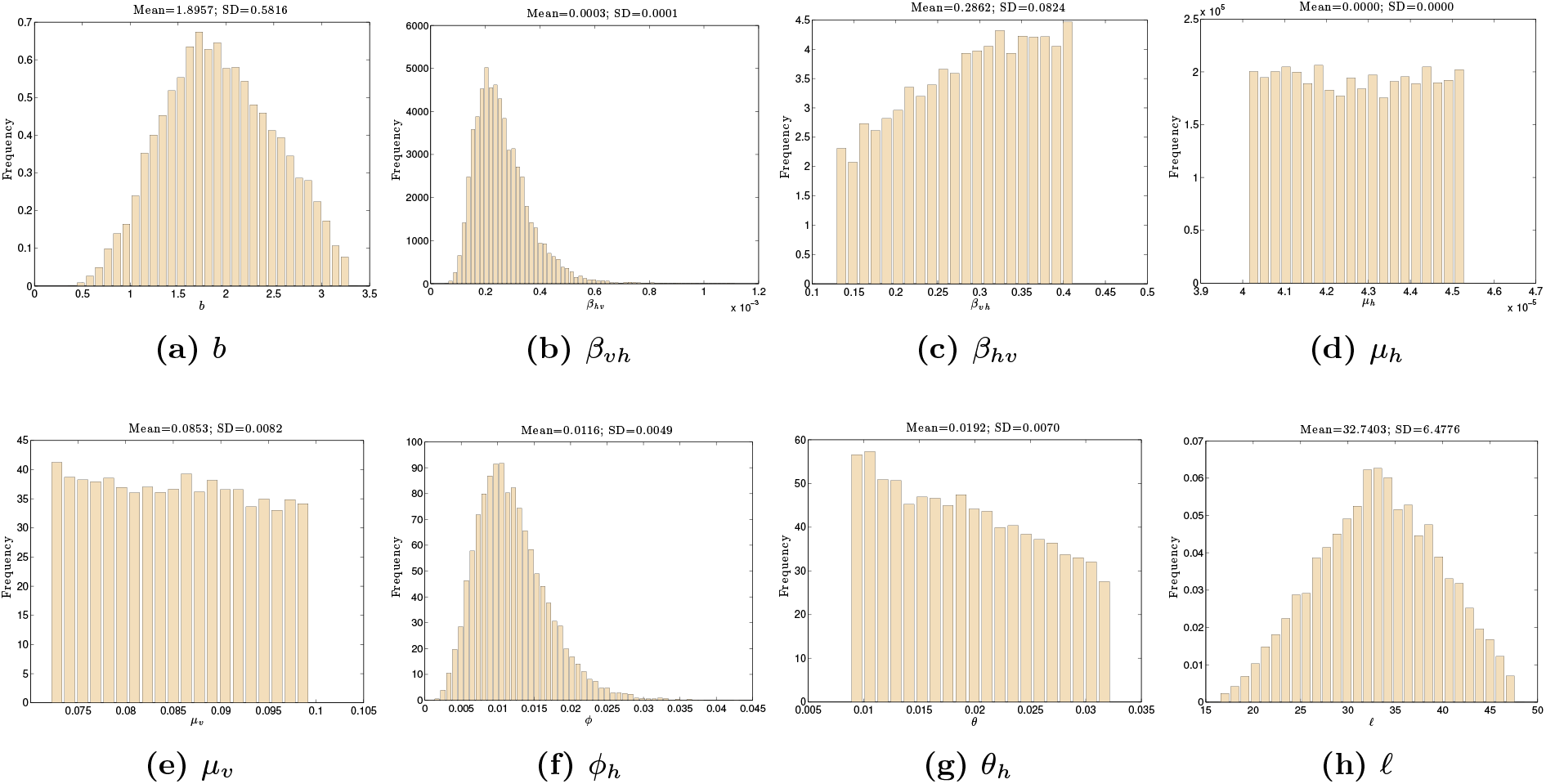
Estimated distributions of the model parameters conditional on 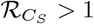 for Sudan obtained from uncertainty analysis of the prevalence

**Figure 9.**
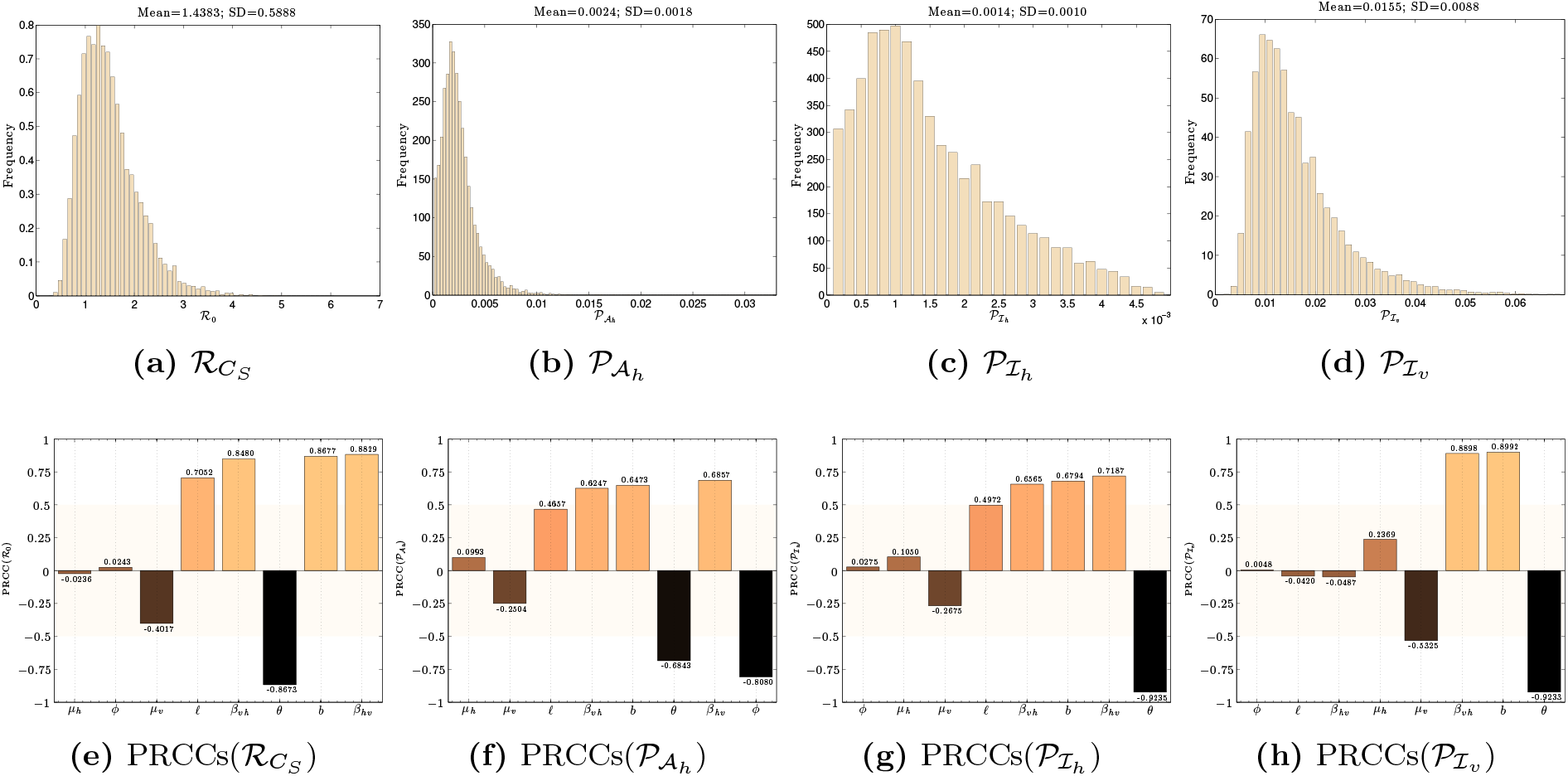
Results for Sudan: Uncertainty of the Reproduction Number (Subfigure 9a) and the Prevalence (Subfigures 9b –9d) of Asymtomatics, Infectious Humans and Infectious Sandflies, respectively. Tornado plot showing partial rank correlation coefficients (PRCCs) of the Reproduction Number (Subfigure 9e) and the Prevalence (Subfigures 9f –9h) of Asymtomatics, Infectious Humans and Infectious Sandflies, respectively.

**Table 8.** The PRCCs by rank of importance for the input parameters of the output values of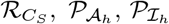, and 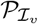 for Sudan. (*) denotes *p* < 0.01.

| Output | $\mathcal{R}_{C_S}$ | | $\mathcal{P}_{A_h}$ | | $\mathcal{P}_{I_h}$ | | $\mathcal{P}_{I_v}$ | |
| --- | --- | --- | --- | --- | --- | --- | --- | --- |
| Rank | Parameter | PRCC | Parameter | PRCC | Parameter | PRCC | Parameter | PRCC |
| 1 | $\beta_{hv}$ | 0.88 | $\phi_h$ | -0.81 | $\theta_h$ | -0.92 | $\theta_h$ | -0.92 |
| 2 | $b$ | 0.87 | $\beta_{hv}$ | 0.69 | $\beta_{hv}$ | 0.72 | $b$ | 0.9 |
| 3 | $\theta_h$ | -0.87 | $\theta_h$ | -0.68 | $b$ | 0.68 | $\beta_{vh}$ | 0.89 |
| 4 | $\beta_{vh}$ | 0.85 | $b$ | 0.65 | $\beta_{vh}$ | 0.66 | $\mu_v$ | -0.53 |
| 5 | $\ell$ | 0.71 | $\beta_{vh}$ | 0.62 | $\ell$ | 0.5 | $\mu_h$ | 0.24 |
| 6 | $\mu_v$ | -0.4 | $\ell$ | 0.47 | $\mu_v$ | -0.27 * | $\beta_{hv}$ | -0.049* |
| 7 | $\phi_h$ | 0.024* | $\mu_v$ | -0.25 * | $\mu_h$ | 0.1* | $\ell$ | -0.042* |
| 8 | $\mu_h$ | -0.024* | $\mu_h$ | 0.099* | $\phi_h$ | 0.028* | $\phi_h$ | 0.0048* |

#### 4.4.2 Uncertainty and Sensitivity Analysis on the Endemic Infected Prevalence

For the asymptomatic prevalence, 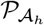, we estimated a mean prevalence of 0.0024 with a standard deviation of 0.0018. Results of uncertainty analysis is shown for Sudan in Figure 9b. From Table 8 and Figure 9f, we observe that the prevalence of asymptomatic population is negatively correlated but most sensitive to *ϕ_h_*, followed by the parameters *β_hv_, θ_h_, b, β_vh_*, and *ℓ*. The natural death rates, *μ_v_*, and *μ_h_*, in humans and sand flies, respectively were the least sensitive input parameters to the prevalence of asymptomatic humans. From our uncertainty analysis on 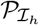 (Figure 9c), we found the average prevalence estimate to be 0.0014 with a standard deviation of 0.0010. The results of our sensitivity analysis, summarize in Table 8 and displayed in Figure 9g shows that the treatment rate of infectious humans, *θ_h_*, is the most influential parameter in determining prevalence level of clinical infection in humans. The infection related parameters, *β_hv_, b, β_vh_* and *ℓ*, also plays a dominant role in disease persistence, but less than *θ_h_*. Finally, the result of uncertainty analysis on the prevalence of infection in sand flies, 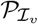, shown in Figure 9d. The estimated sample mean of 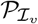 is 0.0155 with a standard deviation of 0.0088. Our analysis identified the parameters sensitivity to changes in 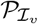 (Table 8 and Figure 9h). The result shows that the treatment rate, *θ_h_* is the most dominant parameter followed by *b, β_vh_*, and *μ_v_*. The less influential parameters on 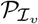 are *μ_h_, ℓ, ϕ_h_*, and *β_vh_*.

### 4.5 Comparative Assessment of VL in India and Sudan

Parameter estimates were obtained either from the literature or estimated from field data, and were used for an evaluation of country-specific risks. The risk was quantified to study differences and similarities in VL disease burden in India and Sudan. In this section, we conduct comparative (between two countries) assessment by studying impact of change in parameter estimations on VL disease burden in these two countries when risk is measured either in terms of 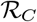 or prevalence of infection. The assessment was based on uncertainty and sensitivity analyses..

#### 4.5.1 Comparsion when risk is defined based on reproduction number

The observed difference in the mean estimate of 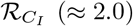 and 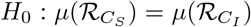 could be because India has much higher levels of endemicity (almost more than 40%) as compared to Sudan. Statistical test was carried out to identify if there exist any significant differences in the estimated means of 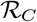 for India and Sudan (t-test with 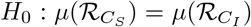 against 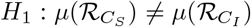 where the *μ* represents mean of 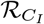 and 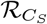). The analysis suggested rejection of null hypothesis (Table 9), that is, the obtained point estimates of 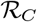 between India and Sudan are different. We also performed Kolmogorov-Smirnov test between empirical distributions of 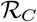 for the two countries and found that empirical distributions are not the same.

**Table 9.**
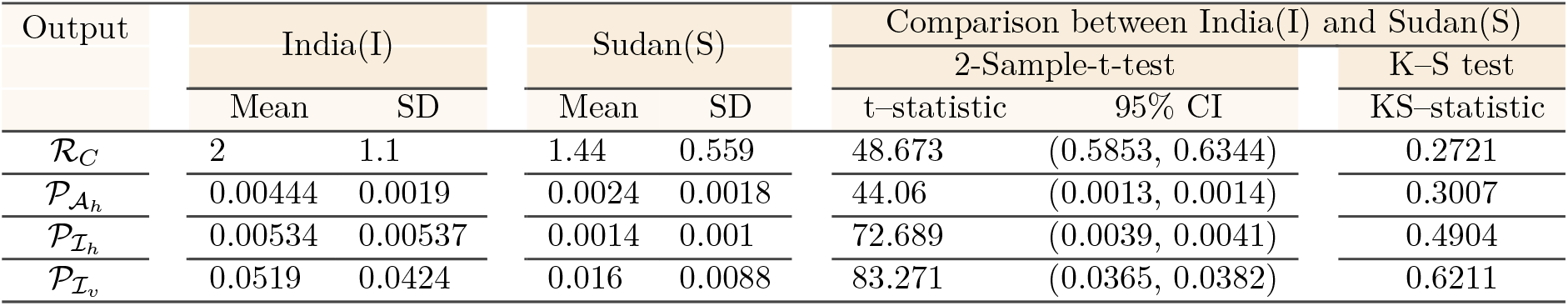
Statistical estimates of quantities,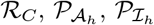 and 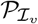, for VL in Sudan and India using the 2 sample t-test and two-sample Kolmogorov–Smirnov test. All analysis were found to be significant, i.e. *p* < 0.05.

| Output | India(I) |  | Sudan(S) |  | Comparison between India(I) and Sudan(S) |  |  |
| --- | --- | --- | --- | --- | --- | --- | --- |
|  | Mean | SD | Mean | SD | 2-Sample-t-test |  | K-S test |
|  |  |  |  |  | t-statistic | 95% CI | KS-statistic |
| $\mathcal{R}_C$ | 2 | 1.1 | 1.44 | 0.559 | 48.673 | (0.5853, 0.6344) | 0.2721 |
| $\mathcal{P}_{A_h}$ | 0.00444 | 0.0019 | 0.0024 | 0.0018 | 44.06 | (0.0013, 0.0014) | 0.3007 |
| $\mathcal{P}_{I_h}$ | 0.00534 | 0.00537 | 0.0014 | 0.001 | 72.689 | (0.0039, 0.0041) | 0.4904 |
| $\mathcal{P}_{I_v}$ | 0.0519 | 0.0424 | 0.016 | 0.0088 | 83.271 | (0.0365, 0.0382) | 0.6211 |

The outcome of the sensitivity analysis (shown in Table 10 and Figures 12; in order of magnitude) highlights difference in influence of parameters for India and Sudan. In Figures 12 we observe the sign and the magnitude of the PRCC values for each parameter. We observe that all parameter, (namely, *b, ℓ, β_hv_, β_vh_*, and *θ_h_*) are the most important parameters of 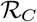 for both countries. The parameters *b, ℓ, β_vh_*, and *β_hv_* with positive PRCC values indicate positive impact on 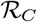 for both countries. The parameter *θ_h_* plays a negative role on the estimation of 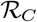, that is, one unit increase in *θ_h_* will result in one unit decrease in 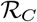 estimate.

**Figure 10.**
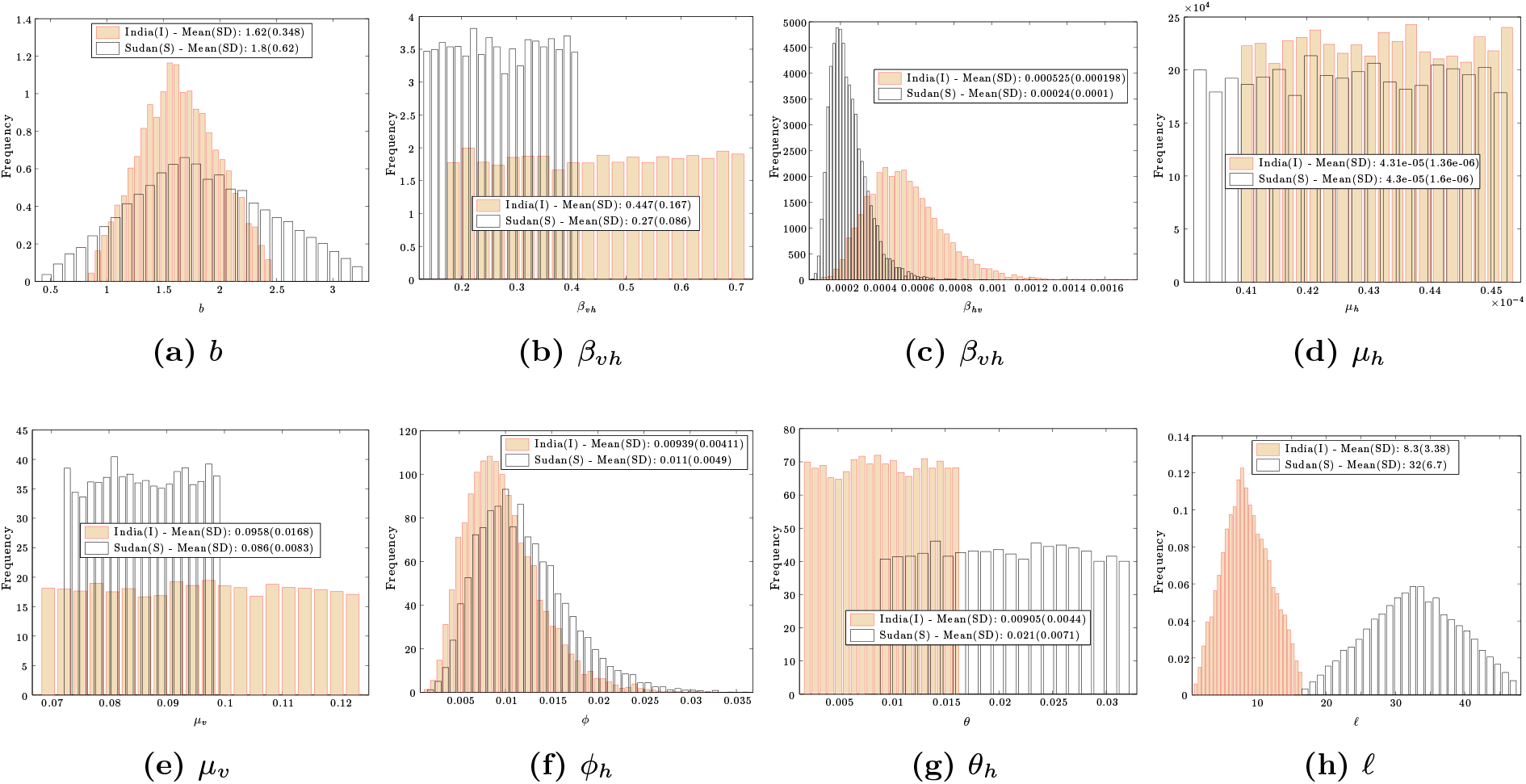
A comparison of initially assigned distributions in Table 6 for model parameters (a) *b*, (b) *β_vh_*, (c) *β_hv_*, (d) *μ_h_*, (e) *μ_v_*, (f) *ϕ_h_*, (g) *θ_h_* and (h) *ℓ* used in the sensitivity and uncertainty analyses for the Indian and Sudan populations

**Figure 11.**
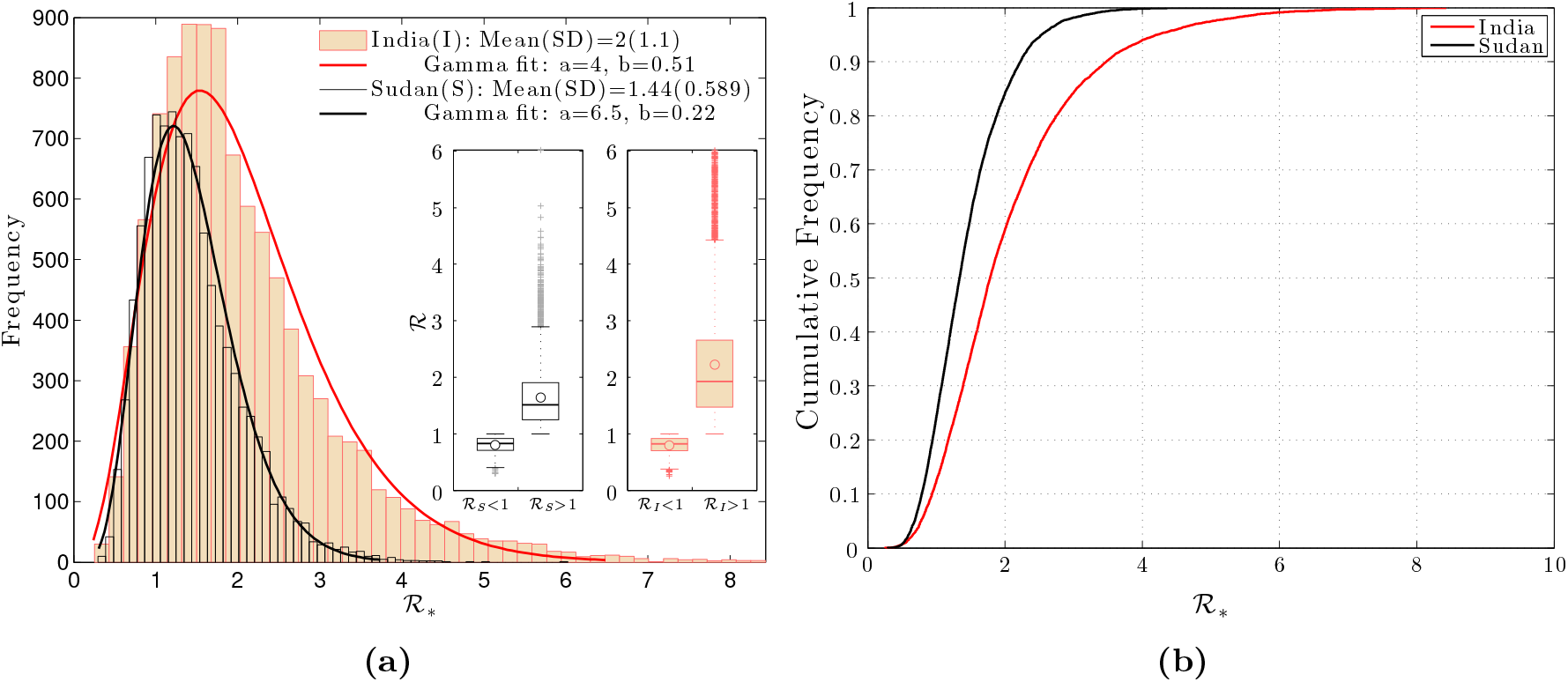
(a) The comparison between estimated distributions of 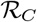 for India and Sudan. The box plot compares the mean(◦), median, minimum, and maximum of 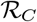 estimates for both countries. It is found that the gamma, is a best-fitted distribution for the samples from the uncertainty analysis. Table 2 summarizes the parameter fitting for the gamma distribution for both countries. (b) The empirical cumulative distributions of the 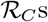 for India and Sudan

**Figure 12.**
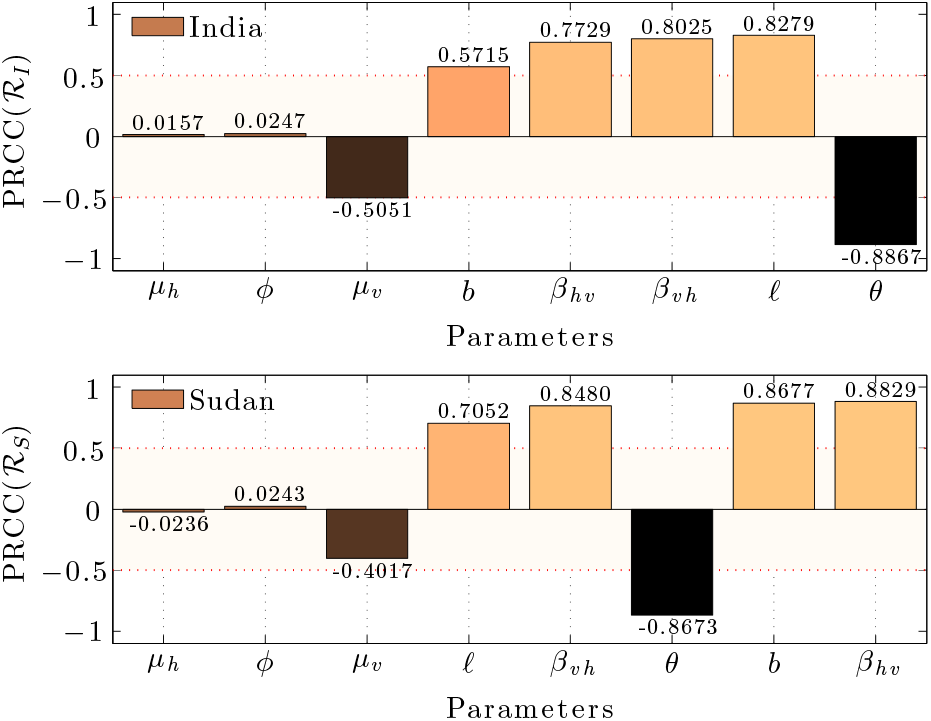
Tornado diagrams of partial rank correlation coefficients, indicating the importance of all eight input parameters that influence the threshold quantity 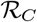. Figure shows a comparison of sensitivity indices for India and Sudan. In both regions, the parameters that have *PRCC* > 0 indicates an increasing influence on 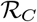 values and those having *PRCC* < 0 will decrease 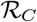 values.

**Table 10.**
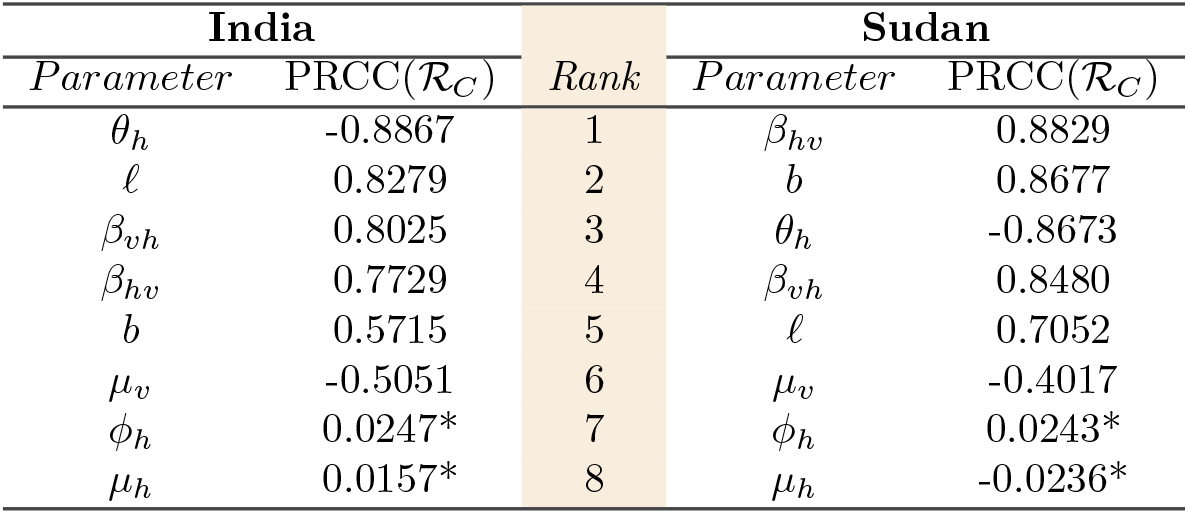
A comparison of the partial rank correlation coefficients for input parameters of the output value 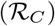, where (*) denotes *p* < 0.01. for India and Sudan.

#### 4.5.2 Comparative assessment if risk is based on different prevalences

##### Point Prevalence of Asymptomatic 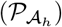

Although the level of 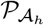, can be determined by how much 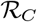 is greater than unity, it is useful to understand the risk posed by an asymptomatic individuals during intensive control. We show that there is a significant difference between the point prevalence of asymptomatic for India (Mean(SD)=0.0037 (0.003)) and Sudan (Mean(SD)=0.0024 (0.002)). There is also a significant statistical difference between the 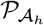-distribution of the two countries (two-sample Kolmogorov–Smirnov test, *p* < 0.050). Combining the results in section 4.3.2 and 4.4.2 we compare the results of sensitivity analysis on 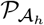 for both countries. We observe from Figure 13c that the most sensitive parameter to both countries in descending order are *ϕ_h_, θ_h_, ℓ, β_vh_, β_hv_*, and *b* and the least sensitive parameter in common to both regions are *μ_v_* and *μ_h_*. From Table 11 and Figure 13c we observed that the two countries differ in order of the parameter ranking with the most sensitive parameter being, *ϕ_h_*. In descending order they are as follows: for India, we have *θ_h_, ℓ, β_vh_, β_hv_*, and *b* and for Sudan, we have *β_vh_, θ_h_, b, β_hv_*, and *ℓ*.

**Figure 13.**
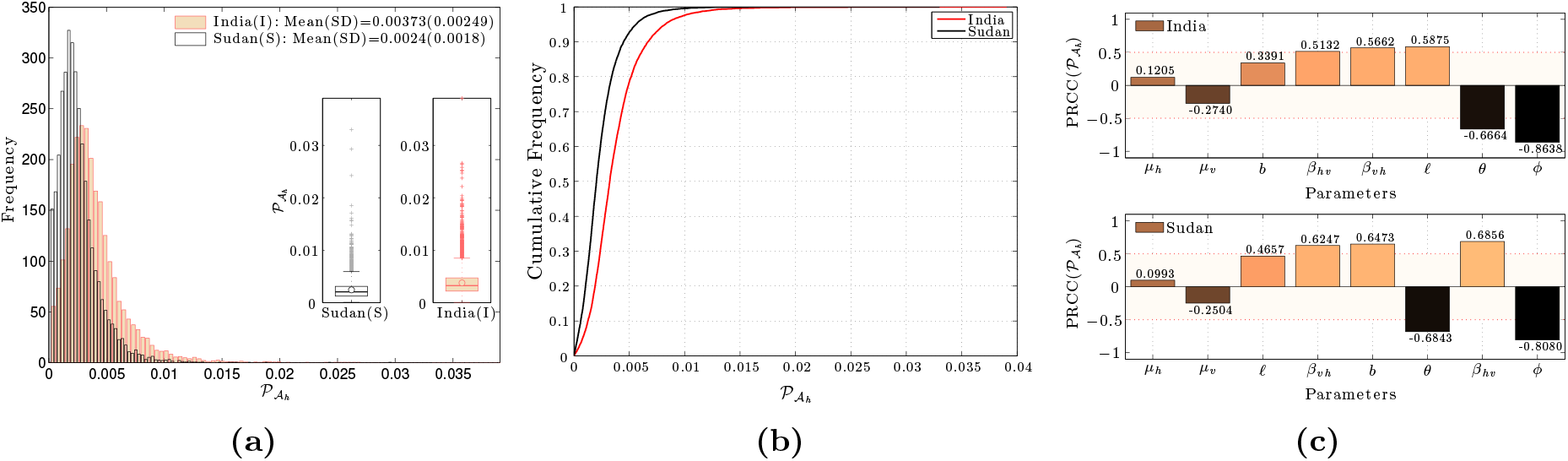
Comparison of uncertainty and sensitivity analysis results on the equilibrium prevalence of asymtomatics humans 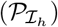: (a) Frequency distributions for contributions, (b) empirical cumulative distributions, and (c) tornado diagrams of partial rank correlation coefficients

**Table 11.** A comparison of the partial rank correlation coefficients for input parameters of the output value 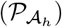. Where (*) denotes *p* < 0.01. for India and Sudan.

| India |  |  | Sudan |  |  |
| --- | --- | --- | --- | --- | --- |
| Parameter | PRCC($\mathcal{P}_{A_h}$) | Rank | Parameter | PRCC($\mathcal{P}_{A_h}$) | |
| $\phi_h$ | -0.8638 | 1 | $\phi_h$ | -0.8080 | |
| $\theta_h$ | -0.6664 | 2 | $\beta_{hv}$ | 0.6856 | |
| $\ell$ | 0.5875 | 3 | $\theta_h$ | -0.6843 | |
| $\beta_{vh}$ | 0.5662 | 4 | $b$ | 0.6473 | |
| $\beta_{hv}$ | 0.5132 | 5 | $\beta_{vh}$ | 0.6247 | |
| $b$ | 0.3391 | 6 | $\ell$ | 0.4657 | |
| $\mu_v$ | -0.2740* | 7 | $\mu_v$ | -0.2504* | |
| $\mu_h$ | 0.1205* | 8 | $\mu_h$ | 0.0993* | |

##### Point Prevalence of Infectious humans 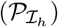

The results showed that there is a significant differences between the point prevalence of Infected humans for India (Mean(SD)=0.0053 (0.005)) and Sudan (Mean(SD)=0.0014 (0.001)). Using p-value< 0.05, the two-sample Kolmogorov–Smirnov test, suggests statistically significant difference between the distributions corresponding to two countries (see Figure 14a - 14b). Sensitivity analysis shows that 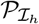 is most sensitive to *θ_h_, ℓ, b, β_vh_*, and *β_hv_* and least sensitive to *μ_h_, μ_v_* and *ϕ_h_* for both countries (Table 12 and Figure 14c). The treatment rate, the first most sensitive parameter, and *μ_v_, μ_h_* and *ϕ_h_* in same decreasing order of influence, are common parameters for both countries. For India, parameters ranking in descending order is *ℓ, β_vh_, β_hv_* and *b* whereas for Sudan the order of parameters is *β_vh_, b, β_hv_*, and *ℓ*.

**Figure 14.**
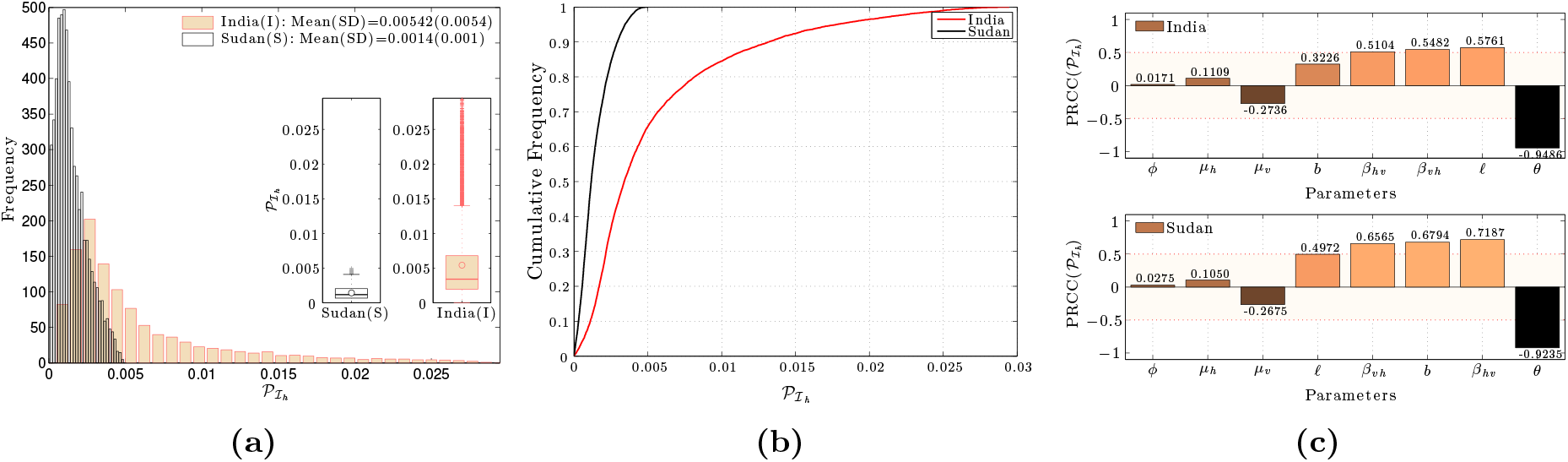
Comparison of result from uncertainty and sensitivity analysis results on the equilibrium prevalence of infected humans 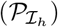: (a) Frequency distributions for contributions, (b) empirical cumulative distributions, and (c) tornado diagrams of partial rank correlation coefficients.

**Table 12.** A comparison of the partial rank correlation coefficients for input parameters of the output value 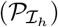. Where (*) denotes *p* < 0.01. for India and Sudan.

| India |  |  | Sudan |  |  |
| --- | --- | --- | --- | --- | --- |
| Parameter | PRCC($\mathcal{P}_{I_h}$) | Rank | Parameter | PRCC($\mathcal{P}_{I_h}$) | |
| $\theta_h$ | -0.9486 | 1 | $\theta_h$ | -0.9235 | |
| $\ell$ | 0.5761 | 2 | $\beta_{hv}$ | 0.7187 | |
| $\beta_{vh}$ | 0.5482 | 3 | $b$ | 0.6794 | |
| $\beta_{hv}$ | 0.5104 | 4 | $\beta_{vh}$ | 0.6565 | |
| $b$ | 0.3226 | 5 | $\ell$ | 0.4972 | |
| $\mu_v$ | -0.2736* | 6 | $\mu_v$ | -0.2675* | |
| $\mu_h$ | 0.1109* | 7 | $\mu_h$ | 0.1050* | |
| $\phi_h$ | 0.0171* | 8 | $\phi_h$ | 0.0275* | |

##### Point Prevalence of of Infected sandflies 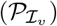

We showed that there is also a significant difference between the point prevalence in infected sand flies for India (Mean(SD)=0.0519 (0.042)) and Sudan (Mean(SD)=0.016 (0.009)), however, there is no statistical difference between the two distributions (Figure 15a - 15b using two-sample Kolmogorov–Smirnov test, *p* < 0.05). Parameters *b, θ_h_, μ_v_*, and *β_vh_* were most sensitive to the prevalence of infection in sand flies, 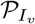, for both countries (see Table 13 and Figure 15c). Between the two parameters, *β_hv_* (b) is relatively more sensitive for India (Sudan). The least important parameter were *μ_h_, ℓ, ϕ_h_*, and *β_hv_* with the exception that the ranks of *ℓ* and *β_hv_* are different.

**Figure 15.**
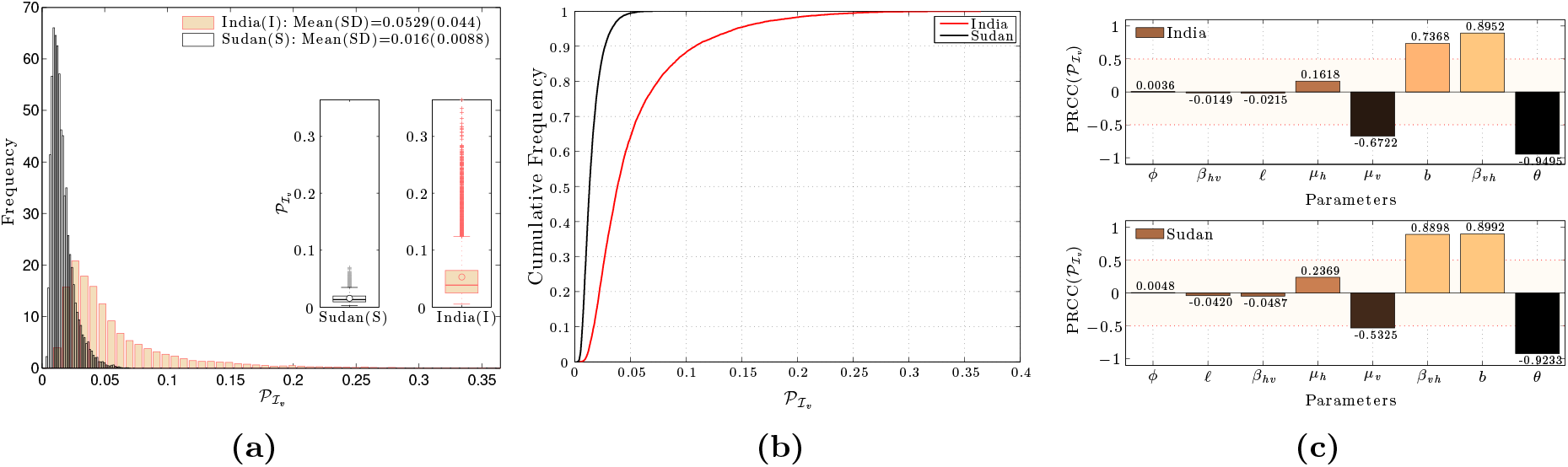
Comparison of uncertainty and sensitivity analysis results on the equilibrium prevalence of infected sandfies 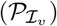: (a) Frequency distributions for contributions, (b) empirical cumulative distributions, and (c) tornado diagrams of partial rank correlation coefficients.

**Table 13.** A comparison of the partial rank correlation coefficients for input parameters of the output value 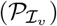. Where (*) denotes *p* < 0.01. for India and Sudan.

| India |  |  | Sudan |  |  |
| --- | --- | --- | --- | --- | --- |
| Parameter | PRCC($P_{I_v}$) | Rank | Parameter | PRCC($P_{I_v}$) | |
| $\theta_h$ | -0.9495 | 1 | $\theta_h$ | -0.9233 | |
| $\beta_{vh}$ | 0.8952 | 2 | $b$ | 0.8992 | |
| $b$ | 0.7368 | 3 | $\beta_{vh}$ | 0.8898 | |
| $\mu_v$ | -0.6722 | 4 | $\mu_v$ | -0.5325 | |
| $\mu_h$ | 0.1618* | 5 | $\mu_h$ | 0.2369* | |
| $\ell$ | -0.0215* | 6 | $\beta_{hv}$ | -0.0487* | |
| $\beta_{hv}$ | -0.0149* | 7 | $\ell$ | -0.0420* | |
| $\phi_h$ | 0.0036* | 8 | $\phi_h$ | 0.0048* | |

## 5 Discussion

The regional risk factors associated with VL are complex and ambiguous. In face of this uncertainty systematic evaluation of ongoing VL control programs is essential but it remains challenging, as appropriate measures of long-term success (where success correspond primarily to no locally acquired cases) with response to changing environmental and political platforms are needed. The objectives of this systematic mathematical analysis is to identify and classify risk factors for India and Sudan using the best available field evidence and data. It will help in determining the gaps in existing knowledge and control and optimally allocating limited resources of the regions. Literature searches were carried out using public health databases, cross sectional and cohort studies, government reports, and information from patients at Rajendra Memorial Institute of Medical Sciences. Due to the limited longitudinal data and no publications with information on comparisons between regions, consistent results could not be found and hence uncertainty and sensitivity analysis help was taken to magnify and identify the missing piece. Most data studies in the literature did not describe information on the criteria of selection of participants in sufficient detail, controlled for confounding variables, or used only one diagnostic test as proof of infection, hence in this study we used multiple data sets to obtained ranges of the parameters.

This is the first study to best of our knowledge that review and make use of extensive collection of available data on epidemiological and ecological parameters to understand the dynamics of the Visceral Leishmaniasis (VL) and identify risk factors in India and Sudan using mathematical modeling approach. The study compares and contrasts quantities from two nations where the disease is endemic and spread via the same VL parasite species and hosts. The sources of the data were used to estimate parameters and uncertainty and sensitivity analysis was conducted on the model’s outcome. Parameter estimates were restricted specifically to India and Sudan to measure the current level of endemicity of VL in both nations. The dynamics of the model depends on the VL basic and control reproductive numbers 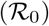 and 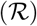, which measures the likelihood and severity of an outbreak. The estimated value of the VL control reproductive number is found to be twice for India(2.1) as compared with Sudan(1.3). Uncertainty analysis on the 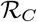 also showed that there were eight parameters (see Table 2) that should be taken into consideration when assessing the uncertainty associated with the risk of increasing levels of VL. The parameter sensitivity analysis 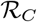 suggests that the biting rate, the average number of vectors per person in a given day, the probability of infection transmission between vector and humans, and the treatment rate were the most influential parameters in the complex disease transmission cycle between sand flies and humans for both countries. However, the order of parameter sensitivity differ between India and Sudan. The biting rate, of the *P. Argentipes* in India and the *P. Orientalis* in Sudan were also shown to be the highest contributing factor to the disease’s severity. Hence, controls reducing the biting rate may be the most effective in controlling VL.

In India the *P. Argentipes* is the main sand fly species responsible for transmission of VL to human populations. During1960’s the man-biting rate of sand flies was significantly reduced from DDT spraying applications that were employed in the malaria eradication campaign and designed to kill mosquito vectors. This campaign reduced the number of VL cases during this period (1962-1963,) showing no new prorated cases. It was observed that soon after the DTT spraying campaign stopped the number of VL cases were elevated to higher epidemic levels [5]. High treatment rate is also found to be a critical factor in impacting the dynamics of VL but primarily in India. However, we assumed effective treatment for all individuals in the model and did not consider efficacy and toxicity of available drugs. These assumptions may influence our findings.

This is an attempt to understand the collective impact of some risk factors contributing to the VL burden in two distinct geographical regions. The results are based on model’s parameter estimates collected and estimated from the available VL data reports. The study was limited to the particular regions of interest as well as to the time period in which data are obtained to estimate some of the parameter estimates from literature were established. As with similar studies, this research also had some limitations. For instance the data used came from various sub-regions and during different time periods and therefore may not be a representative of the country. However, the study clearly identifies the type of data that are relevant and needs to be collected for thoroughly understanding of VL dynamics. In our future research, we plan to provide elaborate analytical methods for the estimation of partially observed data (usually temporal incidence data) for the two developing countries.

## Author Contributions

### Acknowledgements

The study is supported by grants from the National Science Foundation (NSF - Grant DMS-1263374 and 0838705), the National Security Agency (NSA - Grant H98230-13-1-0261 and the Office of the Provost of Arizona State University. The first author would like to thank Ridouan Bani for assistance in programming as well as Sherry Towers and Gerardo Chowell for their helpful suggestions and recommendations on parameter estimation.

## Supporting information

### A Complete Model Derivation

The dynamics of *Leishmania donovani* transmission in humans and sandflies are modeled by the system of equations given by model (1)–(2) in which the force of infection is modeled by Equation C. Newly infected but not yet infectious individuals move into the asymptomatic population (sub-clinical infection, exposed to VL but not yet infectious), who may exit the system through natural death or through progress to clinical VL. The change in *A_h_* population is

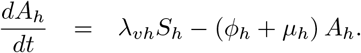

The asymptomatic can then progress to a VL clinical symptoms stage (*I_h_*) at the rate *ϕ_h_*:

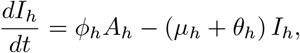

where *θ_h_* is the per-capita treatment rate and *μ_h_* is the per-capita departure rate. The infectious individuals with clinical symptoms may enter treatment (*T_h_*) at the rate *θ_h_*. Through successful treatment, individuals recover at the rate *γ_h_*, and hence

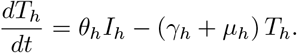

The population of recovered individuals from VL (*R_h_*) is increased following successful treatment, leading to permanent immunity into the *R_h_* class (at the rate *γ*_h_). The population is decreased by natural death and is given by

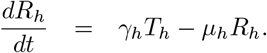

The population of new female sandflies (*S_v_*) is increased by an adult recruitment rate (*λ_v_*) and decrease by natural mortality (*μ_v_*). The vector in this population can acquire the *L. Donovani parasite* from an infectious human at a rate *λ_v_* and is modeled by Equation 4. The change in the susceptible population is described by

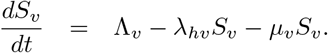

The population of infected female sandflies is generated at the per-capita rate *λ_hv_* and diminished by the natural death rate *μ_v_*. Thus,

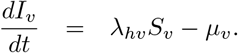

### B Details of the Analytical Results of VL Model

#### B.1 Derivation of the Control Reproductive Number

For simplification, we let *G*_1_ = *ϕ_h_* + *μ_h_,G*_2_ = *θ_h_* + *μ_h_* and *G*_3_ = *γ_h_* + *μ_h_*. Considering the infected subpopulations *I_h_*(*t*), *A_h_*(*t*), and *I_v_*(*t*), we let 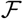 be the rate of new infections into the infected compartments and 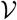 be the rate of exit of humans into infected compartments:

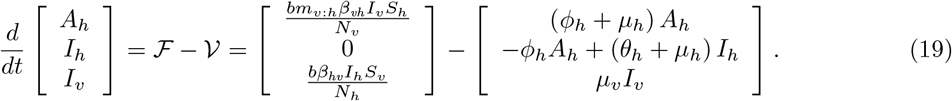

We apply the next generation operator method presented in [84], where 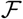 is considered to be the vector of rates of inflow of new infections in each compartment and 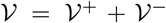 is the vector of rates transfer rates of individuals into and out of the infective compartments by all other processes. Taking the Jacobian matrix of each vector with respect to each of the infectious classes and evaluating at *E*_0_ = (Λ*_h_/μ_h_*, 0, 0, 0, 0, Λ*_v_/μ_v_*, 0) gives

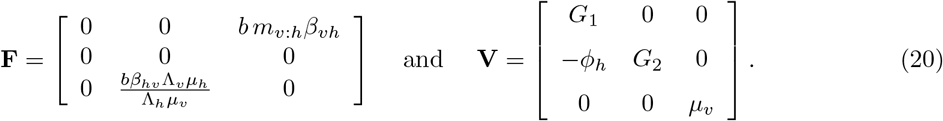

Computing **FV**^−1^, we obtain

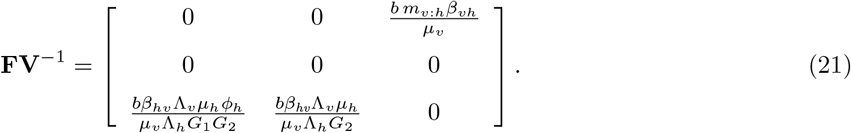

Taking the spectral radius of the next generation matrix operator, *ρ*(**FV**^−1^), gives

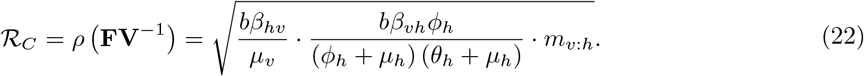

#### B.2 Positivity and Boundedness of Solutions

Since this model is of epidemiological relevance, all its associated parameters are non-negative. Further, the following non-negativity result holds. The state variables of the model (1) are non-negative for all time, so solutions are positively invariant in Ω = Ω_*h*_ × Ω_*v*_, where

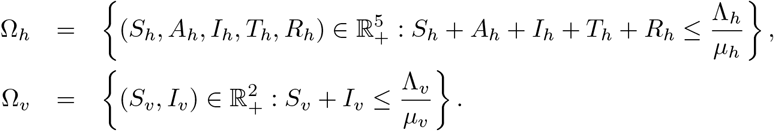

*Remark* B.1. If all initial conditions start in region Ω = Ω_*h*_ × Ω_*v*_, then all corresponding solutions (*S_h_, A_h_, I_h_, T_h_, R_h_, S_v_, I_v_*)′ are non-negative for all *t* > 0, where ′ means vector transpose.

##### Proof.

Because this model is of epidemiological relevance, we first show that the region Ω is positively invariant in 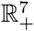, with respect to the system (1) and (2). It is easy to see that 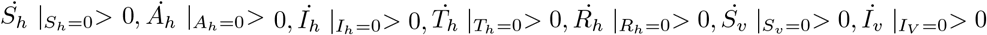. Hence, all trajectories point to inside the region Ω (where the dot means derivative with respect to time). Also, the time derivative along all solutions of (1) is

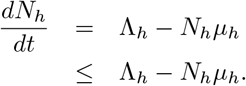

It is clear that *dN_h_/dt* < 0 if *N_h_* > Λ_*h*_/*μ_h_*. Hence, on applying a (comparison) theorem from Birkhoff and Rota ([10]) on differential inequality, we get

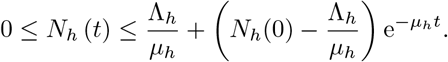

When *t* → ∞, then *N_h_* < Λ_*h*_/*μ_h_*. Thus, for initial conditions *N_h_*(0) < Λ_*h*_/*μ_h_*, we have *N_h_*(*t*) < Λ_*h*_/*μ_h_*.

Similarly, let 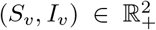 be the solution with non-negative initial solution. Taking the time derivative along the sum of all solutions curves of model (2) gives

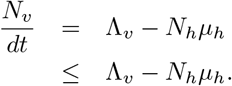

By differential inequality theorem in [10], we find

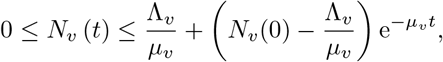

where *N_v_*(0) represents the initial sandfly population at the initial phase of the disease. As *t* → ∞, the inequality becomes

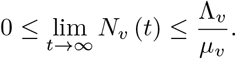

In particular, we have *N_v_*(*t*) < Λ_*v*_/*μ_v_* if *N_v_*(0) < Λ_*v*_/*μ_v_*. Hence the region Ω is positively invariant. Furthermore, if we start with initial conditions *N_h_* (0) > Λ_*h*_/*μ_h_* and *N_v_*(0) > Λ_*v*_/*μ_v_*, then either the solutions enter Ω in finite time or *N_h_*(*t*) → Λ_*h*_/*μ_h_* and *N_v_*(*t*) → Λ_*v*_/*μ_v_*, as *t* → ∞.

Hence, for the model (1–2), the compact set Ω is a positively invariant and absorbing set that attracts all solutions of model (1–2) starting in 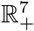.

#### B.3 Stability Analysis of the Disease-Free Equilibrium Point (DFE)

##### B.3.1 Local stability of the Endemic Equilibrium (DFE)

*Remark* B.2. The disease-free equilibrium point, *E*_0_, of model system 1-2 is locally asymptotically stable (LAS) if 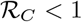, and unstable if 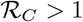.

###### Proof.

Linearization at DFE gives

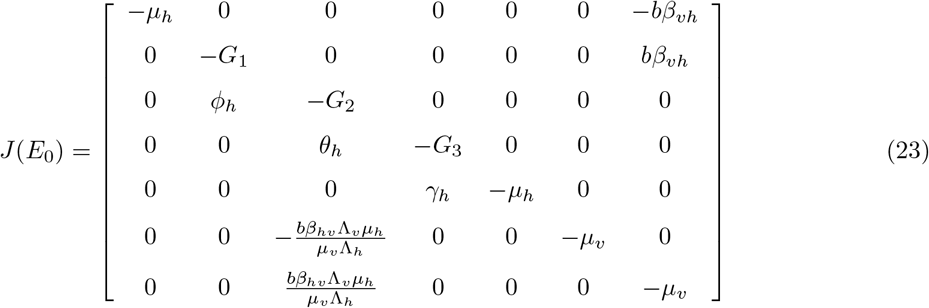

The characteristic polynomial of the Jacobian matrix *J*(*E*_0_) is given by

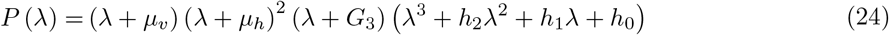

where 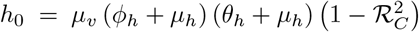, *h*_1_ = (*G*_1_ + *G*_2_)*μ_v_* + *G*_1_*G*_2_ and *h*_2_ = *G*_2_ + *G*_1_ + *μ_v_*. We observe that four eigenvalues for this polynomial have negative real parts, and are given by λ = {−*μ_v_*, −*G*_3_, −*μ_h_*, −*μ_h_*} with geometric multiplicity of two. The remaining expression is a cubic polynomial, *P*(λ) = λ^3^ + *h*_2_λ^2^ + *h*_1_λ + *h*_0_. Applying the Routh-Hurwitz criteria [43], we find the conditions for all eigenvalues to have negative real parts, that is *H*_1_ = *h*_1_ > 0, *H*_2_ = *h*_0_ > 0, and *H*_3_ = *h*_2_*h*_1_ − *h*_0_ > 0. Thus by Routh-Hurwitz criteria, *E*_0_ is locally asymptotically stable for 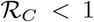 and is unstable for 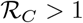.

##### B.3.2 Global Stability of the Disease-free Equilibrium (DFE)

*Remark* B.3. The disease-free equilibrium 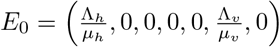 of model system 1-2 is globally asymptotically stable in Ω whenever 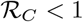 and unstable if 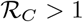.

###### Proof.

Consider a candidate Lyapunov function defined in Ω,

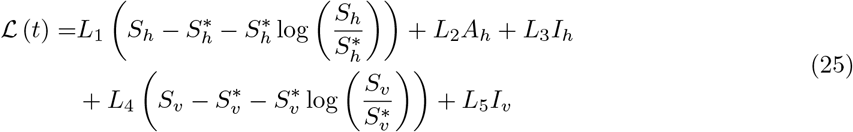

where the constants *L_i_,i* = 1…5 are taken to be 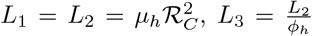, and 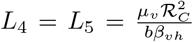. The function 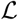 is positive definite, in the sense that it vanishes only at the disease-free equilibrium while otherwise it is positive in Ω. Moreover, taking the time derivative of the function in (25) along solutions of system 1–2 and then substituting the expression for the derivatives, gives

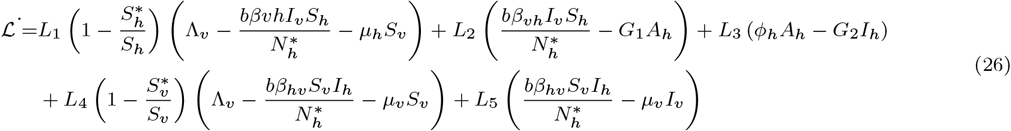

Substituting the *L_i_* constants in equation 26 and then grouping and collecting terms, gives

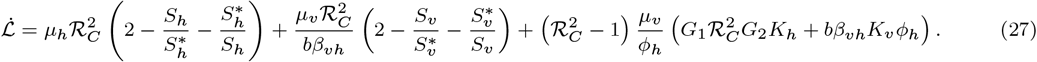

The first two terms are negative, as the arithmetic mean is greater than or equal to the geometrical mean. However, the third term is negative for values of 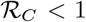. Therefore, by Lyapunov-LaSalle asymptotic stability [52], the disease-free equilibrium *E*_0_ is globally asymptotically stable if 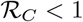 for all *t* > 0.

#### B.4 Stability Analysis of the Endemic Equilibrium Point, *E*\*

As a result of no disease deaths, observeD in Figure ??, the existence of a DFE and an Endemic Equilibrium (EE) that depends on 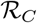. In this section, we show the local and global stability of the EE when 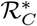 become 1.

*Remark* B.4. If 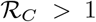, then the unique positive endemic equilibrium(EE), *E*\*, for Model system (1–2) is locally asymptotically stable.

##### Proof.

The EE of the Model system equations 1–2 is given by *E*\*. The Jacobian matrix at EE gives by

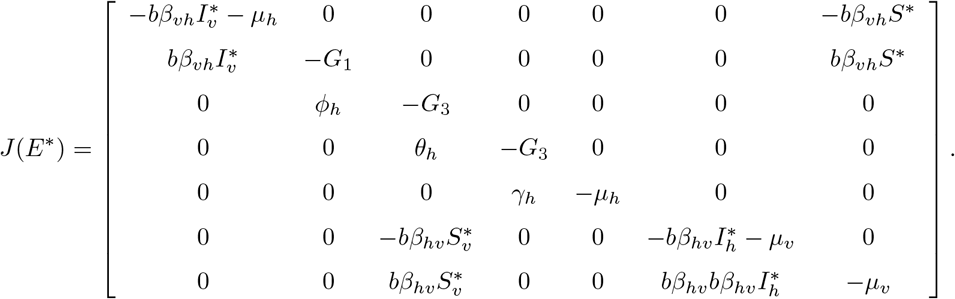

It’s characteristic polynomial is given by

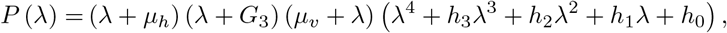

where

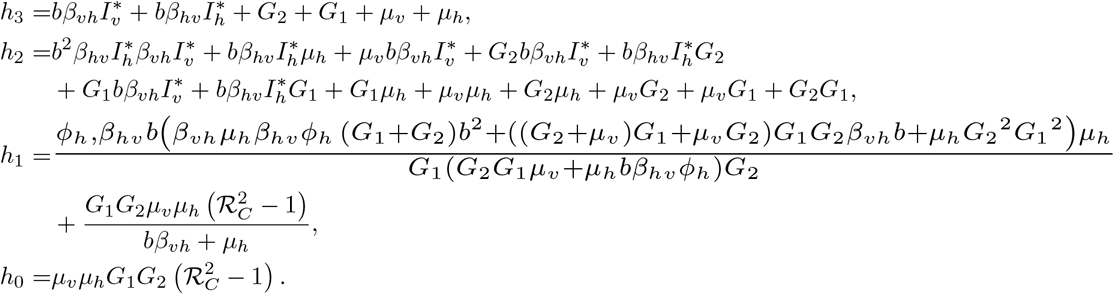

We observe that the characteristic polynomial *P*(*λ*) can be factored to roots *λ* = −*μ_h_*, −*μ_v_*, −*G*_3_ and 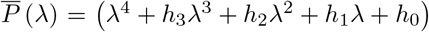. Applying the Routh-Hurwitz conditions: *h_i_* > 0, (*i* = 0,…, 4), *h*_1_*h*_2_ − *h*_0_*h*_3_ > 0, and 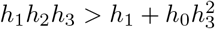, we find that

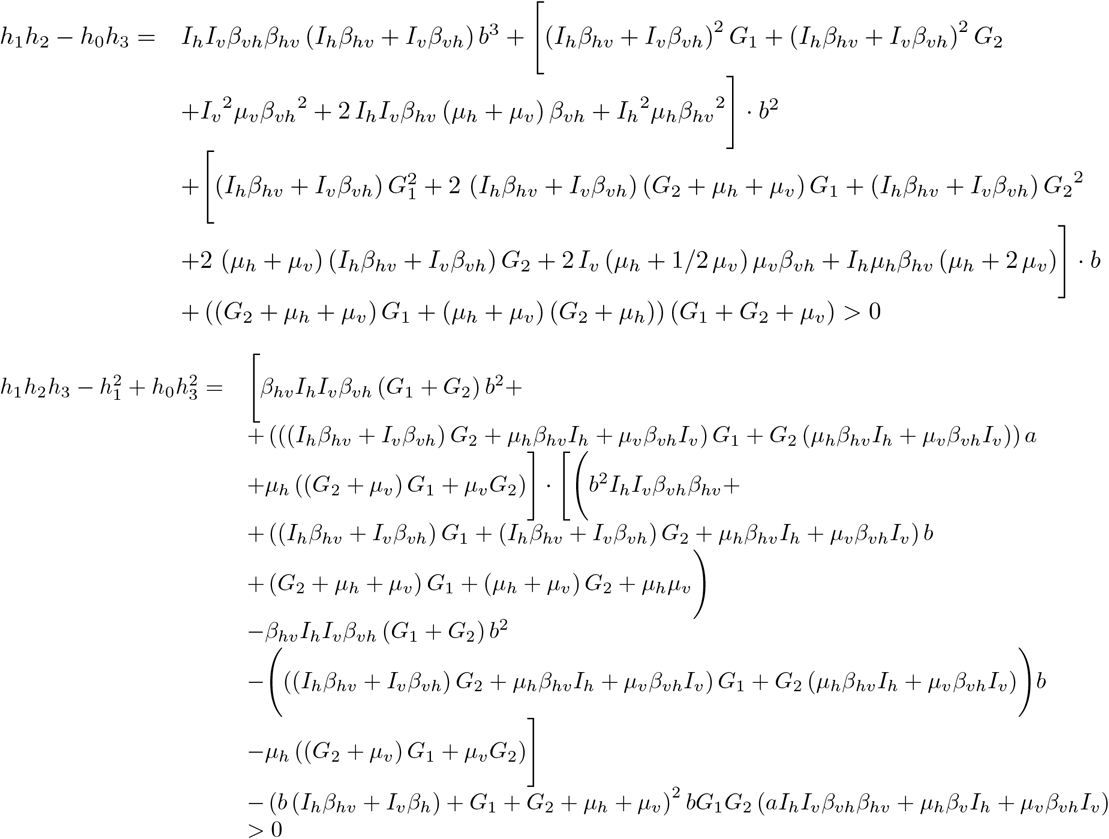

hold when 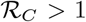. Thus, the endemic equilibrium, *E*\*, is locally asymptotically stable because all eigenvalues of the septic polynomial have all negative real parts for 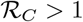.

##### B.4.1 Global stability of the Endemic Equilibrium (EE)

*Remark* B.5. If 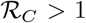, then the unique positive endemic equilibrium, *E*\*, for Model (1–2) is globally asymptotically stable.

###### Proof.

Consider a candidate Lyapunov function defined in Ω,

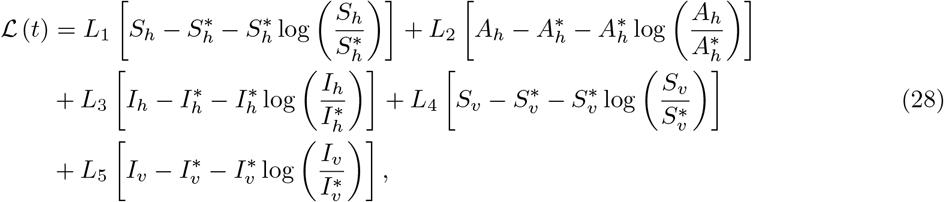

where the constants *L_i_, i* = 1…5 are given by 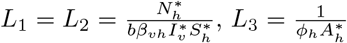, and 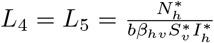. Taking the time derivative of the Lyapunov function in (28) along solutions of system 1–2 and then substituting the expression for the derivatives gives

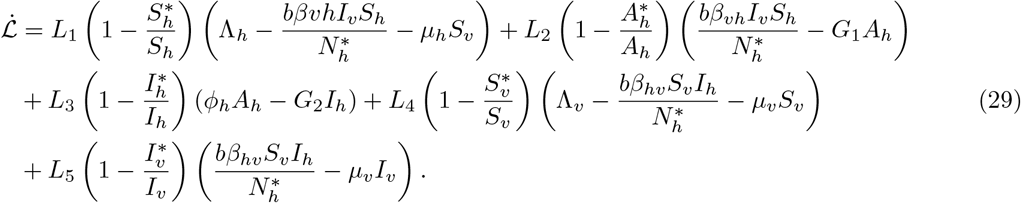

Substituting the *L_i_* in 29 and performing some algebra gives

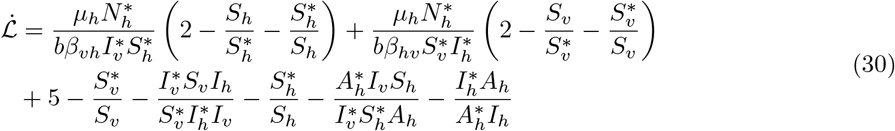

The first two terms in parenthesis and the remaining expression are negative, as the arithmetic mean is greater than or equal to the geometrical mean. Therefore, by LaSalle’s Invariable Principle [52], the endemic equilibrium point *E*\* is globally asymptotically stable in Ω for 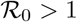 for all *t* > 0.

### C Estimating Model Parameters

After extensive searching of the literature, annual reports, and census data, ecological and epidemiological parameter ranges for the respective human and sandfly populations in India and Sudan were gathered and estimated. See Table 2 for a summary of these estimates.

*b*: The per-capita daily biting rate on humans by female Phlebotomus sandflies species differ by geographical region.

**P. Argentipes (India):** On average, the biting rate of a sandfly on a human per night was estimated to be 0.85 *per day* and range from 0.2 to 2.5 per day [51]. More current studies found a mean estimates biting density per day to be 0.7997 with a range of 0.1667 to 2.0833 per day [22]. From these studies, we calculated the mean number of bites on a human to be 0.7997 with a range of 0.1667 to 2.083 bites per human.
**P. Orientalis (Sudan):** In a field investigations conducted by Elnaiem, et al., the average bites *per man-night* was estimated to range from 23.7 to 40.3 for no bed net and 4.2 to 9.6 for those using untreated bed nets over a period of 12 nights [27]. In both studies, an average of 32 bites per *man-night* was established over a period of 12 nights. In our model we took the average biting rate to be 1.6208 per man-night with a range of 0.35 to 3.3583 per man-night.

*β_hv_*: The transmission probability that an uninfected sandfly acquires a VL parasite from an infectious human.

**India** Parameter estimates were taken from a recent modeling study on VL in India by Stauch A, et al. [77, 78]. From these, we took the mean transmission potential to be 0.025 with a range between 0.013 and 0.063.
**Sudan** We use the infection rate for sandflies, using an equation from our model to estimate *β_hv_*. We first solve for *β_hv_* in this expression and use average infection rates of 9.6% [72], 8.6% [39] and 6.9%, and 3.6% [30] and the average biting rates in Table 2. The average transmission potential in human for *P. Orientalis* was estimated to be 0.1275 with a range of 0.0640 to 0.1706.

*β_vh_:* The transmission probability, is the probability that a VL-infectious sandfly transmits to a human.

**India** Parameter estimates were generated by solving for *β_vh_* in our 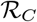 expression

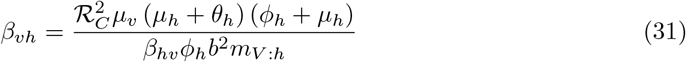

and then pairing samples of known values in Table 2 together with an estimated 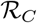 value of 2.01 by Mubayi, et al. (2010 [57]). From this calculation, the mean transmission coefficients were estimated as 0.0694 with a range of 0.0266–0.1652.
**Sudan** A similar approach from India was taken and applied to Sudan using know parameter estimates from Table 2 and an estimated 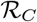 value of 1.3 from ELmojtaba, et al, 2010 [26]. The calculations yield an average estimate for *β_vh_* as 0.0012 with a range of 0.0007–0.0020.

*μ_v_*: The per-capita daily mortality rate of an adult sandfly, taken as 1/ (life expectancy of sandflies)

**P. Argentipes (India)** The mortality for this species of sandfly varies between 0.125 to 0.1 [75] and 0.0667 to 0.1 [65] per day. Some studies established the average lifespan to be, 0.0833 per day [44,50] and 0.091 per day [77]. For this species, the per-capita mortality rate was averaged out from these studies to be *μ_v_* = 0.0833 per day with a range of 0.0667 to 0.1 per day.
**P. Orientalis (Sudan)** The adult life span of this species has not been well studied. In one extensive study, the whole life cycle range was 48–60 days [38]. From this study, the combine time of the four (4) different developmental larval stages and the pupation stage gives a range of 40 to 56 days. So, the life span of adult sandflies ranges from 10 to 14 days and average 12 days. For this species, the per-capita mortality rate was averaged out to be, *μ_v_* = 0.0857 per day and ranges from 0.1 to 0.0714 per day.

*ℓ*: The human landing rate of an adult female sandflies was used as a approximate measure of the human biting rate. Before the late 1990s, the human landing catches (HLC), was a common way for measuring the human landing rate of Phlebotomine sandflies. However, for ethical reasons, this method is less commonly used and has been replaced with the use of human baits and Centers for Disease Control light traps (CDCLT) to attract female sandflies. In a comparison study, Dilger, E. (2013) investigated the relationship between the number of sandflies caught by HLC and CDCLT upon humans and showed that CDCLT are appropriate for estimating the number of sandflies visiting humans [21]. Various comparatives on HLC and CDCLT were used as measured to establish an appropriate parameter range for the human landing rate.

**P. Argentipes (India)** In this study conducted by Joshi B, et al. (2009) [45] on the collection of *P. Argentines* per house per night using CDC LT, we took the mean number of landing 12.15 with a range of 8.68 to17.
**P. Orientalis (Sudan)** From a studies conducted on the effectiveness of impregnated bed net on the landing/bite of female *P. Orientalis* human volunteers by Elnaiem et al. (1999, 2011), we took the mean number of human landing rate to be 32 landing/human/per day with a range 15.7 to 48.3 landing/human/per day [27, 32].

*μ*_h_: For both India and Sudan, the average life expectancy at birth in a year was collected from multiple censored data sources. Using these sources, we estimate the per-person/day natural death rate as (average life expectancy ×365)^−1^. For each of these respective regions, the mean and range of the natural death rates was estimated to be:

**India** From the mean data from multiple survey sites, we found the per-capita natural death rate to be 4.55e-5 (Census of India, 2001), 4.28e-5 (hetv.org, 2012), 4.08e-5 (cia.gov, 2010), 4.33e-5 (WHO, 2012), and 4.27e-5 (un.org, 2012). Combining the estimates of these various value gave a mean death rate of per human/day and range of 4.05e-5 to 5.03e-5 per human/day.
**Sudan** Similarly from India, the per-capita natural death rate was found to be 4.55e-5 (Coutinho, 2005), 4.38e – 5 (cia.gov, 2012), 4.49e – 5 (unicef.org, 2012), 4.09e – 5 (WHO, 2012) and 4.54e-5 (un.org, 2012). The mean death rate of 4.3e – 5 per human/day and range of 4.e – 5 to 4.54e – 5 per human/day.

*ϕ_h_*: The per-capita rate of progression of humans from the asymptomatic state to the infectious state here is taken at incubation of VL before becoming symptomatic. The incubating period is known to vary from weeks to years among different individuals.

**India** The day^−1^ asymptomatic rate has been estimated to be 0.0086 (day^−1^) [79], 0.0055 [60,83] and range between 0.0055 – –0.0164 (day^−1^) [17] and 0.0167 to 0.0083 (day^−1^) [63]. We consider these estimates and took the asymptomatic rate incubating period, *ϕ_h_*, to be 0.00975 (day^−1^) with a range of 0.006–0.0167 (day^−1^).
**Sudan** For this region, the day^−1^ asymptomatic rates ranges were estimated to be 0.0083 to 0.01667 (day^−1^) [34], 0.0055 to 0.0164 (day^−1^) [14, 17], and specific mean rates are give in 0.0167 (day^−1^) with a rang of 0.0111 to 0.0042 (day^−1^) [37]. The asymptomatic rate incubating period, taken as an average of all these studies was taken to be *ϕ_h_* = 0.0098 (day^−1^) and range from 0.0042 to 0.0167 (day^−1^).

*θ_h_*: Treatment rate from VL here is defined as the mean duration of illness before seeking treatment in some treatment fertility.

**India** Current estimates for treatment were found to be 1.996 (who2007), 4 months (0.5–19 months) [2], 4 months [7], and 3.5 [8]. From these study we took the mean estimated treatment rate per day was *θ_h_* = 0.0351 (day^−1^) with a range of 0.0067 to 0.0597 (day^−1^).
**Sudan** The estimated mean rates per person/day varied from 0.0164 [18,59], 0.0130, 0.0055 [3], 0.0108 (0.0027–0.0408) [46], (0.0033–0.0235) [56] and a range of 0.0111–0.0056 in [73]. We took the mean estimate for *θ_h_* as 0.014275 (day^−1^) with a range of 0.0027 to 0.0408 (day^−1^).

Λ_*h*_: The per-capita recruitment rates is defined as the sum per-capita birth rate and per-capita net migration rate of the population.

**India** To estimate the per-capita recruitment rate, we use demographic data on population size, birth rate, and migration from CIA World Factbook. The average estimated recruitment rate was calculated as the sum of the birth rate and net immigration per day and is given by 8.3e-5 persons per day, ranging from 7.67e-5 to 9.22e-5 persons per day.
**Sudan** Similar to the estimation for India, the average estimated recruitment was 1.27e-4 persons per day, with a range of 1.1e-4 to 1.35e-4 persons per day.

Λ_*v*_: The per-capita daily adult sandfly recruitment rate of female phlebotomus sandfly. Seasonality plays a role in the abundance of the sandfly population in each geographical region. Few studies have established an average recruitment rate for sandflies to 0.02128 × *N_h_* per day [75] and 0.299 per day [47]. For our model, we consider the recruitment rate for both species to be Λ_*v*_ = 0.1601 per day and range from 0.0213 to 0.299 per day.

*P_I_h__*: Prevalence for VL in humans is defined as the proportion of people with the disease at a given point in time.

**India** To estimate the per day prevalence, a study based on Serodiagnostic Test in Madhepura District of Bihar, India, was considered by Srivastava N, et al., 2014 [76]. From this study, we use the annual prevalence per 10000 of 26.92 in 2010 and 23.78 in 2011 together with the total population of Madhepura assumed to be at risk to estimate the per person per day prevalence. The prevalence range was estimated to be between 0.0013 to 0.0015 persons per day.
**Sudan** A Survey study by Khalil et al. 2000 [48], gave the prevalence of active disease a range from 40 to 80 per 1000. Using these estimates, together with reported estimates of the at risk population in Pigott et al., 2014 [66], a rough estimate of the daily prevalence range of 0.0006 to 0.0013 persons per was generated for Sudan’s population.

*P_I_v__*: Prevalence for VL in sandflies is defined as the proportion of sandflies with VL at a given point in time.

**Table 14.** Point Prevalence Estimates for VL in India and Sudan for Host and Vector From Various Sample-Based Field Studies.

| Species | Point Prevalence of sandflies |  |  |  |
| --- | --- | --- | --- | --- |
|  | Min | Max | Mean | Reference |
| P. argentipes | 0.0085 | 0.0284 | - | [82] |
|  | 0.005 | 0.05 | - | [58] |
|  | 0.007 | 0.02 | - | [64, 69, 71, 81] |
| P. orientalis | 0.019 | 0.05 | - | [32] |
|  | 0.0054 | 0.037 | 0.0157 | [40] |
|  | 0.035 | 0.071 | - | [30] |

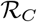: Estimated ranges for both countries were taken from previous mathematical and modeling studies.

**India** 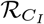 was estimated to 2.0 ± 0.25 [57, 77]
**Sudan** 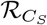 was estimated to be 1.3 ± 0.25 [26]

**Table 15.**
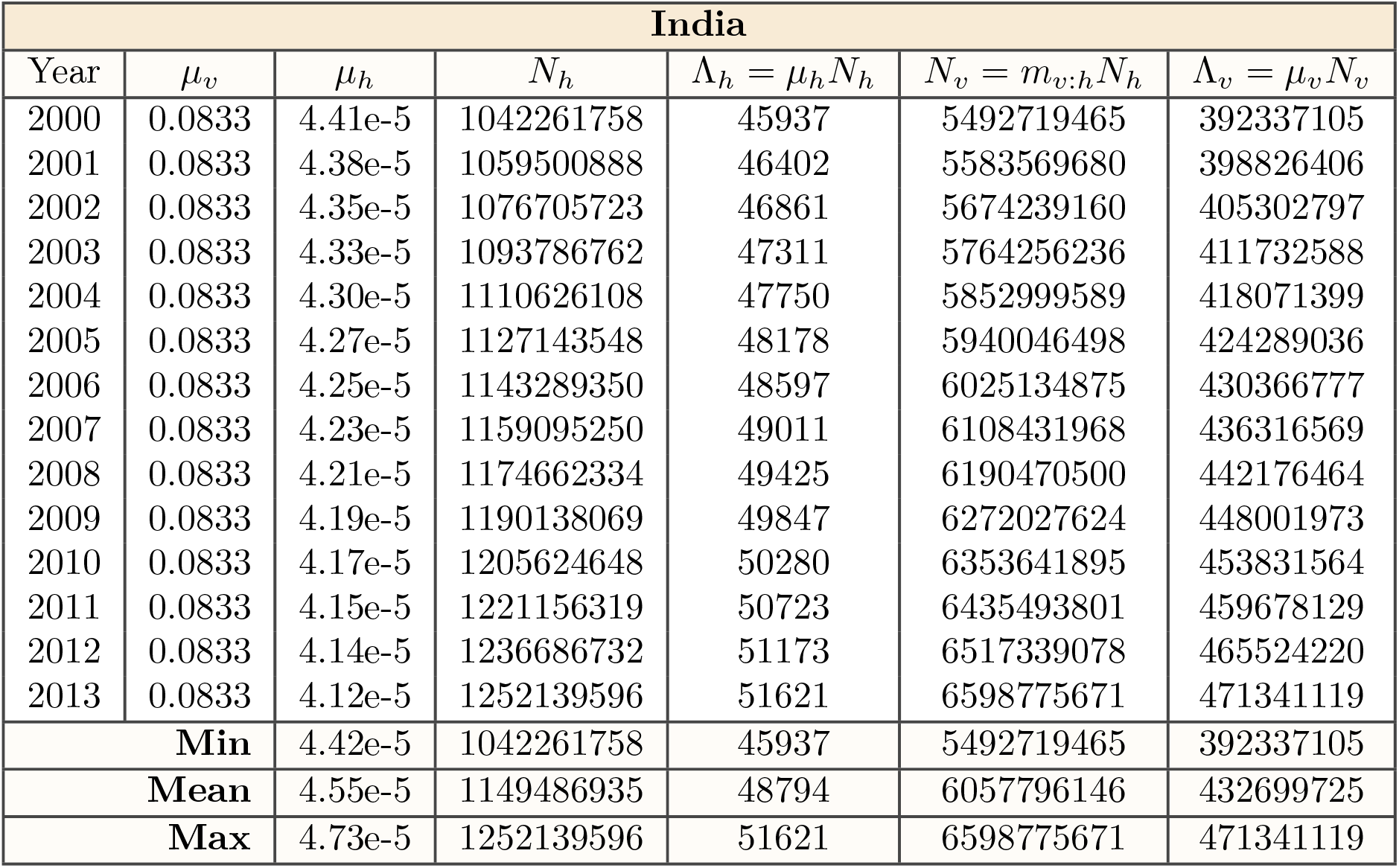
Estimate for Parameters Λ_*h*_ and Λ_*h*_ Using Mean Estimates for India in Table 2 and World Bank’s Demographic Estimates in [35]

| India |  |  |  |  |  |  |
| --- | --- | --- | --- | --- | --- | --- |
| Year | $\mu_v$ | $\mu_h$ | $N_h$ | $\Lambda_h = \mu_h N_h$ | $N_v = m_{v:h} N_h$ | $\Lambda_v = \mu_v N_v$ |
| 2000 | 0.0833 | 4.41e-5 | 1042261758 | 45937 | 5492719465 | 392337105 |
| 2001 | 0.0833 | 4.38e-5 | 1059500888 | 46402 | 5583569680 | 398826406 |
| 2002 | 0.0833 | 4.35e-5 | 1076705723 | 46861 | 5674239160 | 405302797 |
| 2003 | 0.0833 | 4.33e-5 | 1093786762 | 47311 | 5764256236 | 411732588 |
| 2004 | 0.0833 | 4.30e-5 | 1110626108 | 47750 | 5852999589 | 418071399 |
| 2005 | 0.0833 | 4.27e-5 | 1127143548 | 48178 | 5940046498 | 424289036 |
| 2006 | 0.0833 | 4.25e-5 | 1143289350 | 48597 | 6025134875 | 430366777 |
| 2007 | 0.0833 | 4.23e-5 | 1159095250 | 49011 | 6108431968 | 436316569 |
| 2008 | 0.0833 | 4.21e-5 | 1174662334 | 49425 | 6190470500 | 442176464 |
| 2009 | 0.0833 | 4.19e-5 | 1190138069 | 49847 | 6272027624 | 448001973 |
| 2010 | 0.0833 | 4.17e-5 | 1205624648 | 50280 | 6353641895 | 453831564 |
| 2011 | 0.0833 | 4.15e-5 | 1221156319 | 50723 | 6435493801 | 459678129 |
| 2012 | 0.0833 | 4.14e-5 | 1236686732 | 51173 | 6517339078 | 465524220 |
| 2013 | 0.0833 | 4.12e-5 | 1252139596 | 51621 | 6598775671 | 471341119 |
| **Min** |  | 4.42e-5 | 1042261758 | 45937 | 5492719465 | 392337105 |
| **Mean** |  | 4.55e-5 | 1149486935 | 48794 | 6057796146 | 432699725 |
| **Max** |  | 4.73e-5 | 1252139596 | 51621 | 6598775671 | 471341119 |

**Table 16.** Estimate for Parameters Λ_*h*_ and Λ_*h*_ Using Mean Estimates for Sudan in Table 2 and World Bank’s Demographic Estimates in [36]

| Sudan |  |  |  |  |  |  |
| --- | --- | --- | --- | --- | --- | --- |
| Year | $\mu_v$ | $\mu_h$ | $N_h$ | $\Lambda_h = \mu_h N_h$ | $N_v = m_{v:h} N_h$ | $\Lambda_v = \mu_v N_v$ |
| 2000 | 0.0857 | 4.73e-5 | 27729798 | 1310 | 146136035 | 10438288 |
| 2001 | 0.0857 | 4.70e-5 | 28434810 | 1335 | 149851449 | 10703675 |
| 2002 | 0.0857 | 4.67e-5 | 29186427 | 1362 | 153812470 | 10986605 |
| 2003 | 0.0857 | 4.63e-5 | 29973979 | 1389 | 157962869 | 11283062 |
| 2004 | 0.0857 | 4.60e-5 | 30778572 | 1417 | 162203074 | 11585934 |
| 2005 | 0.0857 | 4.57e-5 | 31585871 | 1444 | 166457540 | 11889824 |
| 2006 | 0.0857 | 4.54e-5 | 32397535 | 1472 | 170735009 | 12195358 |
| 2007 | 0.0857 | 4.52e-5 | 33218250 | 1500 | 175060178 | 12504298 |
| 2008 | 0.0857 | 4.49e-5 | 34040065 | 1529 | 179391143 | 12813653 |
| 2009 | 0.0857 | 4.47e-5 | 34853178 | 1559 | 183676248 | 13119732 |
| 2010 | 0.0857 | 4.46e-5 | 35652002 | 1589 | 187886051 | 13420432 |
| 2011 | 0.0857 | 4.44e-5 | 36430923 | 1618 | 191990964 | 13713640 |
| 2012 | 0.0857 | 4.43e-5 | 37195349 | 1647 | 196019489 | 14001392 |
| 2013 | 0.0857 | 4.42e-5 | 37964306 | 1676 | 200071893 | 14290849 |
| **Min** |  | 4.42e-5 | 27729798 | 1310 | 146136035 | 10438288 |
| **Mean** |  | 4.55e-5 | 32817219 | 1489 | 172946744 | 12353339 |
| **Max** |  | 4.73e-5 | 37964306 | 1676 | 200071893 | 14290849 |

